# MEF2C hypofunction in neuronal and neuroimmune populations cooperate to produce MEF2C haploinsufficiency syndrome-like behaviors in mice

**DOI:** 10.1101/824151

**Authors:** Adam J. Harrington, Catherine M. Bridges, Kayla Blankenship, Ahlem Assali, Stefano Berto, Benjamin M. Siemsen, Hannah W. Moore, Jennifer Y. Cho, Evgeny Tsvetkov, Acadia Thielking, Genevieve Konopka, David B. Everman, Michael D. Scofield, Steven A. Skinner, Christopher W. Cowan

## Abstract

Microdeletions of the *MEF2C* gene are linked to a syndromic form of autism termed *MEF2C* haploinsufficiency syndrome (MCHS). Here, we show that MCHS-associated missense mutations cluster in the conserved DNA binding domain and disrupt MEF2C DNA binding. DNA binding-deficient global *Mef2c* heterozygous mice (*Mef2c*-Het) display numerous MCHS-like behaviors, including autism-related behaviors, as well as deficits in cortical excitatory synaptic transmission. We find that hundreds of genes are dysregulated in *Mef2c*-Het cortex, including significant enrichments of autism risk and excitatory neuron genes. In addition, we observe an enrichment of upregulated microglial genes, but not due to neuroinflammation in the *Mef2c*-Het cortex. Importantly, conditional *Mef2c* heterozygosity in forebrain excitatory neurons reproduces a subset of the *Mef2c*-Het phenotypes, while conditional *Mef2c* heterozygosity in microglia reproduces social deficits and repetitive behavior. Together our findings suggest that MEF2C regulates typical brain development and function through multiple cell types, including excitatory neuronal and neuroimmune populations.

## Introduction

Myocyte Enhancer Factor 2 (MEF2) proteins are members of the MADS family of transcription factors that regulate gene expression during development and adulthood. In the brain, MEF2C is important for neuronal differentiation and synapse development (Assali et al., 2019). MEF2 proteins regulate numerous genes associated with synapse formation and function as well as multiple genes linked to neurodevelopmental disorders, including autism spectrum disorder (ASD) (Flavell et al., 2008; Harrington et al., 2016; Morrow et al., 2008). Constitutively-active MEF2C can promote glutamatergic synapse elimination, a process requiring the RNA-binding function of the Fragile X Mental Retardation protein (FMRP) and several other factors (Flavell et al., 2006; Pfeiffer et al., 2010; Tsai et al., 2012; Zang et al., 2013). Conditional knockout of *Mef2c* in neuronal populations within the mouse brain produces a myriad of severe behavioral and synaptic phenotypes, which emphasizes the importance of this gene in healthy brain development (Adachi et al., 2015; Barbosa et al., 2008; Harrington et al., 2016; Li et al., 2008; Rajkovich et al., 2017).

MEF2C in the developing and mature brain is also expressed in microglia (Deczkowska et al., 2017; Gosselin et al., 2017; Zhang et al., 2014a) – a population of macrophage-like cells throughout the brain that regulate synapse formation and pruning during early brain development (Paolicelli et al., 2011; Schafer et al., 2012; Zhan et al., 2014). Microglia influence a number of brain functions, including synapse elimination, synapse formation, fasciculation of the corpus callosum, survival of oligodendrocyte precursor cells, and phagocytosis of other brain cells (Hagemeyer et al., 2017; Parkhurst et al., 2013; Pont-Lezica et al., 2014; Schafer et al., 2012; Shigemoto-Mogami et al., 2014; Sierra et al., 2010; Stevens et al., 2007). Microglia are recognized as not just responding to infection or injury, but as important regulators of brain development and function (Li and Barres, 2018). In addition, microglial dysfunction has been hypothesized to play important roles in disease pathology for other neurodevelopmental disorders, including Rett Syndrome (Derecki et al., 2012; Horiuchi et al., 2017; Schafer et al., 2016; Wang et al., 2015).

Microdeletions on chromosome 5q14.3 that include the *MEF2C* gene or point mutations within the protein-coding region of *MEF2C* are linked to a recently described neurodevelopmental disorder, termed *MEF2C* Haploinsufficiency Syndrome (MCHS) (Berland and Houge, 2010; Bienvenu et al., 2013; Engels et al., 2009; Le Meur et al., 2010; Mikhail et al., 2011; Novara et al., 2010; Paciorkowski et al., 2013; Tonk et al., 2011; Vrecar et al., 2017; Zweier et al., 2010; Zweier and Rauch, 2012). Common symptoms of MCHS include autism spectrum disorder (ASD), absence of speech, stereotypical behaviors, hyperactivity, intellectual disability, hypotonia, motor abnormalities, high pain tolerance, sleep disturbances, and epilepsy, and individuals with *MEF2C* point mutations typically present with fewer and/or milder symptoms (Berland and Houge, 2010; Bienvenu et al., 2013; Engels et al., 2009; Le Meur et al., 2010; Mikhail et al., 2011; Novara et al., 2010; Paciorkowski et al., 2013; Tonk et al., 2011; Vrecar et al., 2017; Zweier et al., 2010; Zweier and Rauch, 2012). Due to the abundance of neurological symptoms and neuronal-enriched expression of MEF2C, *MEF2C* haploinsufficiency in neurons is presumed to underlie most, if not all, of the MCHS symptoms. Interestingly, single-cell genomic profiling from cortical tissue of patients with idiopathic autism revealed that upper-layer excitatory neurons and microglia are preferentially affected in autism (Velmeshev et al., 2019), and since both neurons and microglia express MEF2C, we sought to explore the possible cell type-specific effects of MEF2C hypofunction in MCHS-related behaviors in a construct-valid mouse model of human MCHS.

## Methods and Materials

### Patients

Patients with developmental delay and a significant variant in the MEF2C gene were selected for this study. These patients were seen for clinical genetics evaluations and data from these visits were gathered from records review. Internal informed consent to publish data was obtained for each subject.

### Animals

Mice (*Mus musculus*) were group housed (2-5 mice/cage; unless specified) with same-sex littermates with access to food and water *ad libitum* on a 12-hour reverse light-dark cycle. *Mef2c^+/-^* (*Mef2c*-Het) mice were initially generated by crossing *Mef2c^fl/fl^*(RRID:MGI:3719006) (Arnold et al., 2007) mice with *Prm1-Cre* (Jackson Laboratory #003328) to induce germline recombination of *Mef2c*. *Mef2c^+/-^; Prm1-Cre* were backcrossed with C57BL/6J to remove *Prm1-Cre*, and test mice were generated from *Mef2c^+/-^* mice (male and female) crossed with C57BL/6J mice. *Mef2c* conditional heterozygous mice were generated by crossing *Mef2c* floxed mice with cell type-selective Cre+-expressing transgenic mice (*Emx1-Cre* (Iwasato et al., 2008)), *PV-Cre* (Jackson Laboratory #017320), or *Cx3Cr1^creER/creER^* (Jackson Laboratory #021160 (Parkhurst et al., 2013)) to generate *Mef2c^fl/+^*; *Cre+* conditional heterozygous (*Mef2c* cHet) mice that were compared to their Cre-negative or flox-negative littermates (Control). Experimenters were blinded to the mouse genotype during data acquisition and analysis. Experiments were independently replicated, and the total number of animals/cells were reported in the representative figures. All procedures were conducted in accordance with the Medical University of South Carolina Institutional Animal Care and Use Committee (IACUC) and NIH guidelines.

*Detailed Materials and Methods can be found within the supplemental information*.

## Results

### Patient MEF2C missense mutations cluster in DNA binding and dimerization domains and disrupt DNA binding

Deletions or mutations in *MEF2C* are assumed to create loss-of-function alleles that cause the symptoms of MCHS (Berland and Houge, 2010; Bienvenu et al., 2013; Engels et al., 2009; Le Meur et al., 2010; Mikhail et al., 2011; Novara et al., 2010; Paciorkowski et al., 2013; Tonk et al., 2011; Vrecar et al., 2017; Zweier et al., 2010; Zweier and Rauch, 2012). Given that microdeletions of 5q14.3 often include additional genes beyond *MEF2C*, we sought to identify individuals with mutations within the *MEF2C* protein-coding region, including an intragenic duplication (*i.e.* p.D40_C41dup) and two missense variants (*i.e.* p.K30N and p.I46T) (Table 1). We compared their clinical histories to those associated with two previously reported missense variants in the *MEF2C* gene (Zweier et al., 2010). All five patients presented with global developmental delay and seizures. Seizure onset occurred prior to one year of age in four of the five patients, with the patient carrying the p.I46T variant developing seizures at 21 months. Common features of these individuals included absence of speech, repetitive movements, hypotonia, varied abnormalities on brain MRI, and breathing disturbances. High pain tolerance was noted in two of the patients. There were some minor facial dysmorphisms noted, though there did not seem to be a consistently recognizable gestalt. When a list of additional MCHS mutations was assembled (personal communications), several frameshift and premature stop codon mutations were identified – all of which, if stable, are predicted to produce a truncated MEF2C protein lacking its C-terminal nuclear localization sequence. We also noted that all of the *MEF2C* missense (or small duplication) mutations were clustered within the highly-conserved MADS (DNA binding) or MEF2 (dimerization) domains (Fig. 1A). In a MEF2 response element DNA binding assay, all five of the MADS domain patient mutations caused a loss of MEF2C DNA binding (Fig. 1B-C, S1A), and they did not appear to interfere with wild-type MEF2C DNA binding (Fig. S1B), suggesting a loss-of-function phenotype.

**Table 1:** Summary of Clinical Features of MCHS Patients.

| Table 1. Overview of Clinical Characteristics associated with *MEF2C* variants |  |  |  |  |  |
| --- | --- | --- | --- | --- | --- |
|  | Newly described |  |  | Previously described by Zweier et al. [2010] |  |
| **Demographics** |  |  |  |  |  |
| Sex | F | F | F | F | F |
| Age at evaluation | 13y11m | 17m | 4y3m | 10y5m | 3y |
| **Molecular findings** |  |  |  |  |  |
| *MEF2C* variant | c.120_125dup | c.90G>T | c.137T>C | c.80G>C | c.113T>A |
|  | p.D40_C41dup | p.K30N | p.I46T | p.G27A | p.L38Q |
| Inheritance of variant | Paternal<br>(father mosaic) | de novo | Non-maternal | de novo | de novo |
| **Clinical findings** |  |  |  |  |  |
| Height (percentile) | 23rd | 10-25th | 25th | 25-50th | NR |
| Weight (percentile) | 25th | 25-50th | 40th | 50th | 25th |
| OFC (percentile) | NR | 10-25th | 25th | 50th | 9th |
| Seizures | Yes | Yes | Yes | Yes | Yes |
| Age of onset | 8y | 6m | 21m | 6m | 10m |
| Global developmental delay | Yes | Yes | Yes | Yes | Yes |
| Speech | Few words | Absent | Few words | Absent | Absent |
| Tremors/tremulous | Yes | No | No | NR | NR |
| Hypotonia | Yes (resolved) | Yes | No | Yes | Yes |
| Brain MRI | Abnormal | Abnormal | Normal | Abnormal | Abnormal |
| Abnormalities noted | Areas of heterotopia in left lateral region, subtle but progressive abnormalities of white matter within frontal and parietal regions | Asymmetric appearance of hippocampi, asymmetric enlargement of temporal horn of left lateral ventricle | - | Mild under-myelination of insular cortices bilaterally | Generalized lack of white matter bulk and delay in myelin maturation |
| Repetitive movements | Yes | No | Yes | NR | NR |
| Breathing abnormalities | Yes | Yes | NR | Yes | No |
| Jugular fossa pit on anterior neck | No | No | No | NR | NR |
| High pain tolerance | Yes | NR | Yes | NR | NR |
| Sleeping difficulties | Yes | No | No | NR | NR |
| Social abnormalities | No | Yes | NR | NR | NR |
| Anxiety | No | No | NR | NR | NR |
| Autism spectrum disorder | No | No | Yes | NR | No |
| **Dysmorphic Findings** |  |  |  |  |  |
| Midface | Hypoplastic | Full cheeks | Normal | NR | NR |
| Palpebral fissures | Upslanted | Upslanted with epicanthal folds | Normal | Downslanted | Upslanted |
| Nose | Sharp nasal tip, hypoplastic alae nasi, prominent columella | Normal | Normal | NR | NR |
| Mouth | Small | Bowed upper lip, small mandible | Dental crowding | Dental crowding, full upper lip | Bowed upper lip, widely spaced teeth |
| Chest | Mild pectus excavatum | Normal | Normal | NR | NR |
| Back | Mild lordosis | Normal | Normal | NR | NR |
| Digits | Long, increased space between toes 1 and 2 | Normal | Normal | NR | NR |
| Other | - | Thin hair | - | Thick hair, large ears, fleshy ear lobes | Large ears, prominent ear lobes |

NR=Not Reported

**Figure 1.**
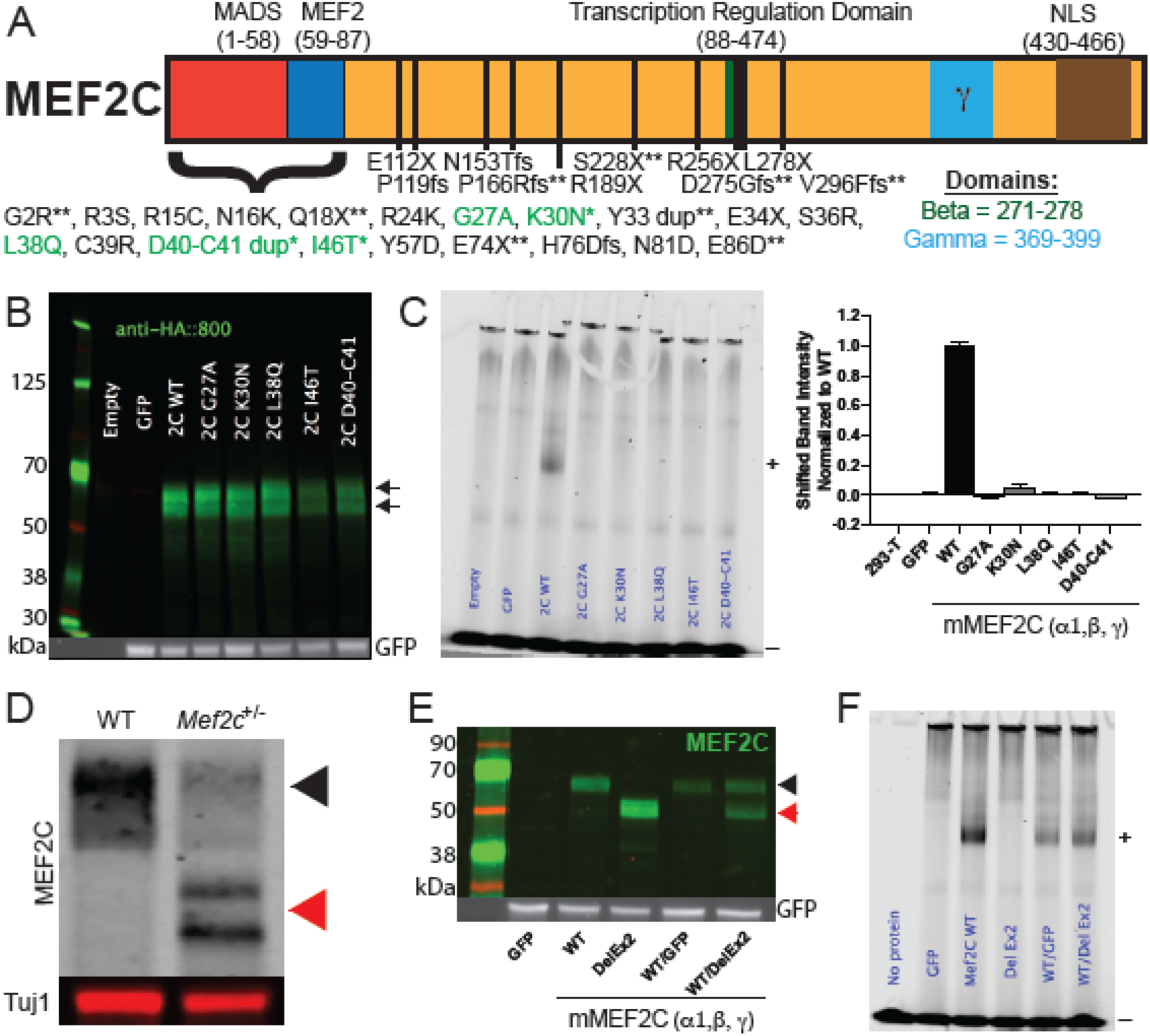
MCHS associated mutations in MEF2C disrupt DNA binding. (A) Schematic of the MEF2C protein with locations of MCHS mutations. MCHS mutations in green are further characterized (B-C). MCHS mutations that are newly described in this manuscript are denoted with “*”. MCHS mutations not previously reported (personal communications) are denoted by “**”. The alternatively spliced beta (green) and gamma (blue) domains are shown. All MEF2C transcripts contain a C-terminal Nuclear Localization Sequence (NLS) that is disrupted by the frame-shift (fs) mutations. (B) Western blot of MEF2C wild-type (WT) and MCHS mutations in 293-T cells show that all MCHS mutations lead to protein expression. Arrows denote WT and mutant protein MEF2C bands. (C) Electrophoretic mobility shift assay (EMSA) using fluorescently labeled MEF2 response element (MRE) probe and MEF2C protein lysates from 293-T cells containing MEF2C mutations. MEF2C bound probe is shifted in the gel (denoted by “+”). Unbound fluorescent probe is denoted with a “–”. Only MEF2C WT binds to the fluorescently labeled MRE, while MCHS mutant proteins fail to bind the MRE probe (C). Quantification of bound probe is included (C); n=3. (D) Western blot of MEF2C from cortical lysates of control and *Mef2c*-Het mice. The black arrow denotes MEF2C WT and red arrows denote MEF2C DelEx2 (D,E). (E) Western blot of MEF2C WT and MEF2C DelEx2 from 293-T cells. (F) MEF2C DelEx2 fails to bind the MRE probe and does not interfere with MEF2C WT binding to MRE probes. “+” is bound probe. “-“ is unbound probe. Data are reported as mean ± SEM. Also see Figure S1.

### Mef2c heterozygous mouse model

To model the genetics of MCHS in mice, we generated a global heterozygous *Mef2c* mutant mouse lacking exon 2 (*Mef2c*^+/^*^Δ^*^Ex2^ or *Mef2c*-Het) (Fig. 1D), which encodes a large portion of the MADS/MEF2 domains. The near full-length MEF2C*^Δ^*^Ex2^ protein had no detectable DNA binding affinity and did not reduce DNA binding affinity of wild-type MEF2C expressed at a similar level (Fig. 1E-F, S1C). We observed a non-Mendelian frequency of *Mef2c*-Hets, suggesting a partial embryonic lethality (Extended Data Fig. 1D, **p<0.01, Chi-square test), similar to a previous report (Tu et al., 2017).

We examined whether male and female *Mef2c*-Het mice showed behavior phenotypes reminiscent of MCHS symptoms. Using a three-chamber social interaction (SI) test, we observed that *Mef2c*-Het mice have a lack of social preference with a novel same-sex mouse (Fig. 2A, *p<0.05, 2-way ANOVA). We also found that *Mef2c-*Het male and female pups (P7-P10) produced significantly fewer ultrasonic vocalization (USV) calls during maternal separation (Fig. 2B, main effect of genotype, **p<0.01), and young adult *Mef2c-*Het males produced significantly fewer USV calls (Fig. 2C, *p<0.05) in the presence of an estrous female, suggesting that *Mef2c-*Hets have deficits in a putative species-appropriate form of social communication. Male *Mef2c-*Hets were hyperactive in a novel environment (Fig. 2E, ***p<0.005) and displayed an increase in jumping (Fig. 2F, *p<0.05), a repetitive-type motor behavior; however, young adult *Mef2c-*Hets displayed normal performance on the accelerating rotarod test of motor coordination (Fig. 2D). In addition, *Mef2c*-Hets showed increased exploration of the open, unprotected arm of the elevated plus maze (Fig. 2G). High pain tolerance is also frequently reported in MCHS individuals (Paciorkowski et al., 2013; Zweier et al., 2010). Interestingly, *Mef2c-*Het mice showed a reduction in startle response to electrical foot-shocks (Fig. 2H; 2-way ANOVA, significant interaction between genotype and shock intensity; *p<0.05, **p<0.01); however, this might be specific to pain stimuli (nociception) given that white-noise acoustic startle responses are indistinguishable from controls (Fig. S2A). Despite a common occurrence of intellectual disability in MCHS, in the *Mef2c*-Hets we failed to detect any clear learning and memory-related deficits in Pavlovian fear conditioning tests (Fig. S2B-D), the Barnes maze test for spatial learning and memory (Fig. S2E), and the Y-maze test for spatial working memory i (Fig. S2F). These mice also showed a strong preference for the novel object in the novel object recognition test (Fig. S2G), and normal sucrose preference in a two-bottle choice test (Fig. S2H). In the cognitively-demanding operant sucrose self-administration (SA) assay, the *Mef2c*-Hets displayed wild-type levels of operant learning, operant discrimination (active vs. inactive port), context-related sucrose seeking after one-week of abstinence, extinction learning and cue-induced reinstatement of sucrose seeking on the formerly-active port (Fig. 2I-J, S2I-L). Taken together, our findings suggest that, unlike the conditional knockout of *Mef2c* in Emx1-lineage cells (Harrington et al., 2016), the global loss of one functional copy of *Mef2c* in mice is not sufficient to produce detectable deficits in learning or memory in the C57BL6/J genetic background.

**Figure 2.**
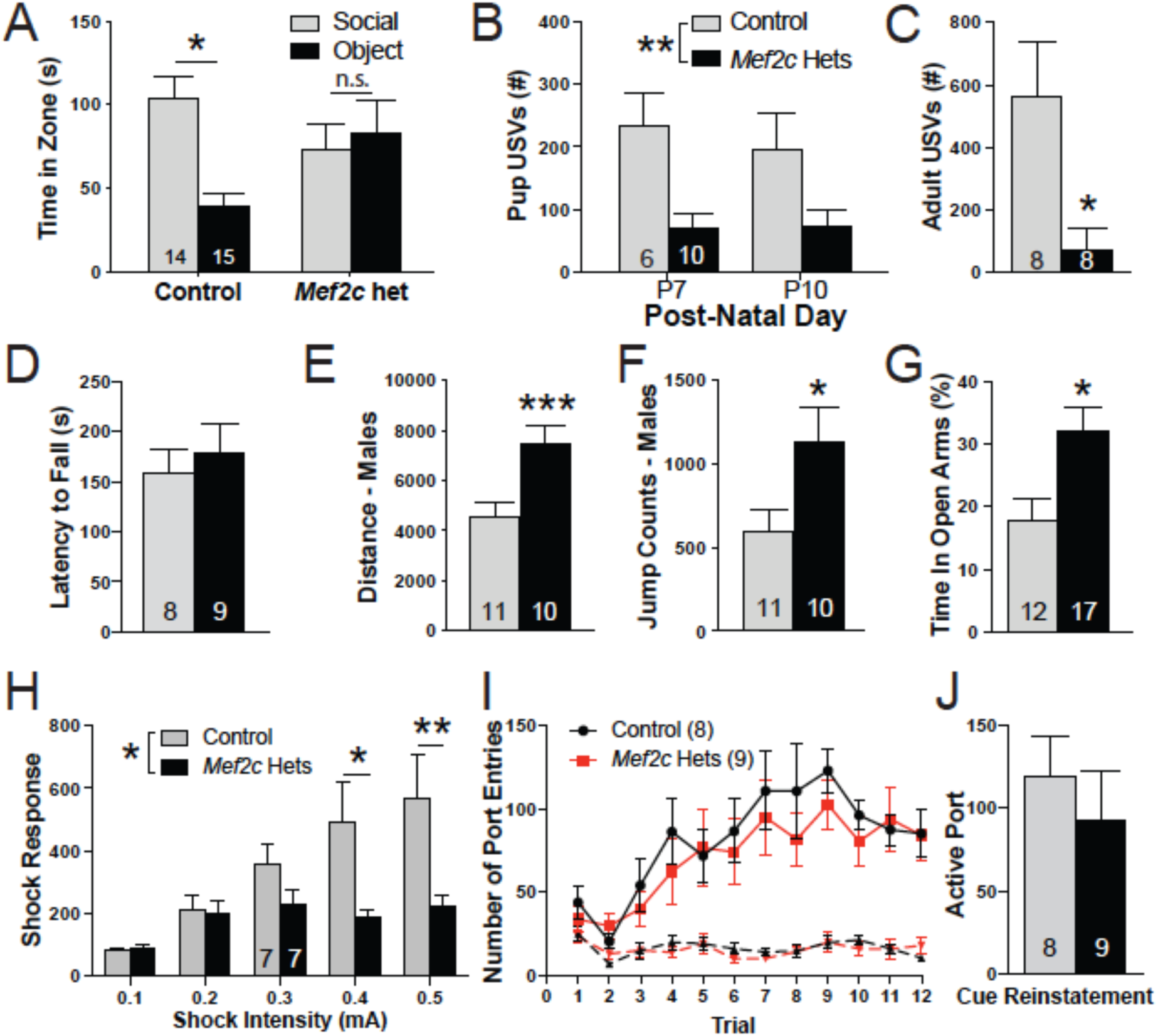
*Mef2c*-Het mice display multiple MCHS-relevant behaviors. (A) Three chamber social interaction test. Control mice spent significantly more time interacting with a novel animal over a novel object while the *Mef2c*-Het mice showed no preference for the novel object or the novel animal. (B) *Mef2c*-Het pups emitted fewer ultrasonic vocalizations (USVs) during maternal separation in early post-natal development. (C) Adult male *Mef2c*-Het mice produced fewer USVs that control mice in the presence of a female mouse in estrous. (D) Both control and *Mef2c*-Het mice have similar latencies to fall on an accelerating rotarod. (E,F) Male *Mef2c*-Het mice are hyperactive (E) and show increased jump counts (F). (G) *Mef2c*-Het mice spend significantly more time on the open arms of the elevated-plus maze. (H) *Mef2c*-Het mice have reduced response to shock. (I) Both control and *Mef2c*-Het mice increase the number of active port entries (solid line) during sucrose self-administration. Dashed line represents inactive port entries. (J) Both control and *Mef2c*-Hets show similar active port entries during cue-induced reinstatement of sucrose seeking. Data are reported as mean ± SEM. Statistical significance was determined by 2-way ANOVA (A,B,H,I) or unpaired t-test (C-G,J). *p<0.05, **p<0.01, ***p<0.005, n.s. = not significant. Number of animals (n) are reported in each graph for respective experiment. Also see Figure S2.

### Mef2c-Het mice display input-selective reductions in cortical excitatory synaptic transmission

Changes in excitatory (E) and/or inhibitory (I) synaptic transmission are associated with numerous neuropsychiatric disorders, including ASD (Antoine et al., 2019; Garber, 2007; Zoghbi and Bear, 2012). Conditional knockout of *Mef2c* alters somatosensory cortex (SSCtx) pyramidal neuron E/I synaptic transmission (Harrington et al., 2016; Rajkovich et al., 2017). In the global *Mef2c*-Hets (p35-p40), gross structural organization of barrel fields within cortical layer 4 of the SSCtx appeared normal (Fig. 3A). In SSCtx layer 2/3 pyramidal neurons, we detected no significant differences by genotype for intrinsic excitability (Fig. S3A), dendritic spine density or spine head diameter of apical or basal dendrites (Fig. S3B), or GABA-mediated inhibitory synaptic transmission (mIPSCs; Fig. 3B). However, patch-clamp recordings of layer 2/3 neurons revealed an input-selective deficit in glutamatergic synaptic transmission. Stimulation of horizontal fibers in layer 2/3 of a neighboring cortical column produced a significant reduction in the amplitude of evoked excitatory postsynaptic currents (eEPSCs) (Fig. 3C), suggesting a reduction in pre- and/or postsynaptic transmission. Paired-pulse facilitation (PPF) analysis (50 ms interstimulus interval) of local horizontal inputs revealed a significant increase in PPF ratio (Fig. 3C), indicating a decrease in presynaptic release probability (Fioravante and Regehr, 2011). These effects were input-selective given that electrical stimulation of layer 4 (within the same cortical column) produced eEPSC and PPF responses in layer 2/3 neurons that were indistinguishable from controls (Fig. 3D). To examine if reductions in AMPA-mediated postsynaptic strength might also contribute to the reduced horizontal eEPSCs (Fig. 3C), we measured miniature EPSCs (mEPSCs) under conditions where action potentials are blocked pharmacologically. In layer 2/3 cells from *Mef2c*-Hets, we observed a significant reduction in mEPSCs amplitude (Fig. 3E), suggesting an overall reduction in AMPA-mediated postsynaptic strength. Similar to layer 2/3, we also observed a significant reduction of mEPSC amplitude in SSCtx layer 5 pyramidal neurons of *Mef2c-*Hets (Fig. 3F), suggesting that the reduction in glutamatergic postsynaptic strength is not limited to a specific cortical layer. Consistent with layer 2/3 pyramidal neurons, we did not observe any differences in dendritic spine density or dendritic spine head diameter in basal dendrites from layer 5 pyramidal neurons (Fig. S3C). There was no effect of genotype on layer 5 mEPSC frequency (Fig. 3F), but we observed a significant increase in the layer 2/3 mEPSCs frequency (Fig. 3E) that was not explained by an increase in dendritic spine density (Fig. S3B) or effects on presynaptic functions of local inputs (Fig. 3C,D), and might represent a compensatory effect of long-range connections (Rajkovich et al., 2017).

**Figure 3.**
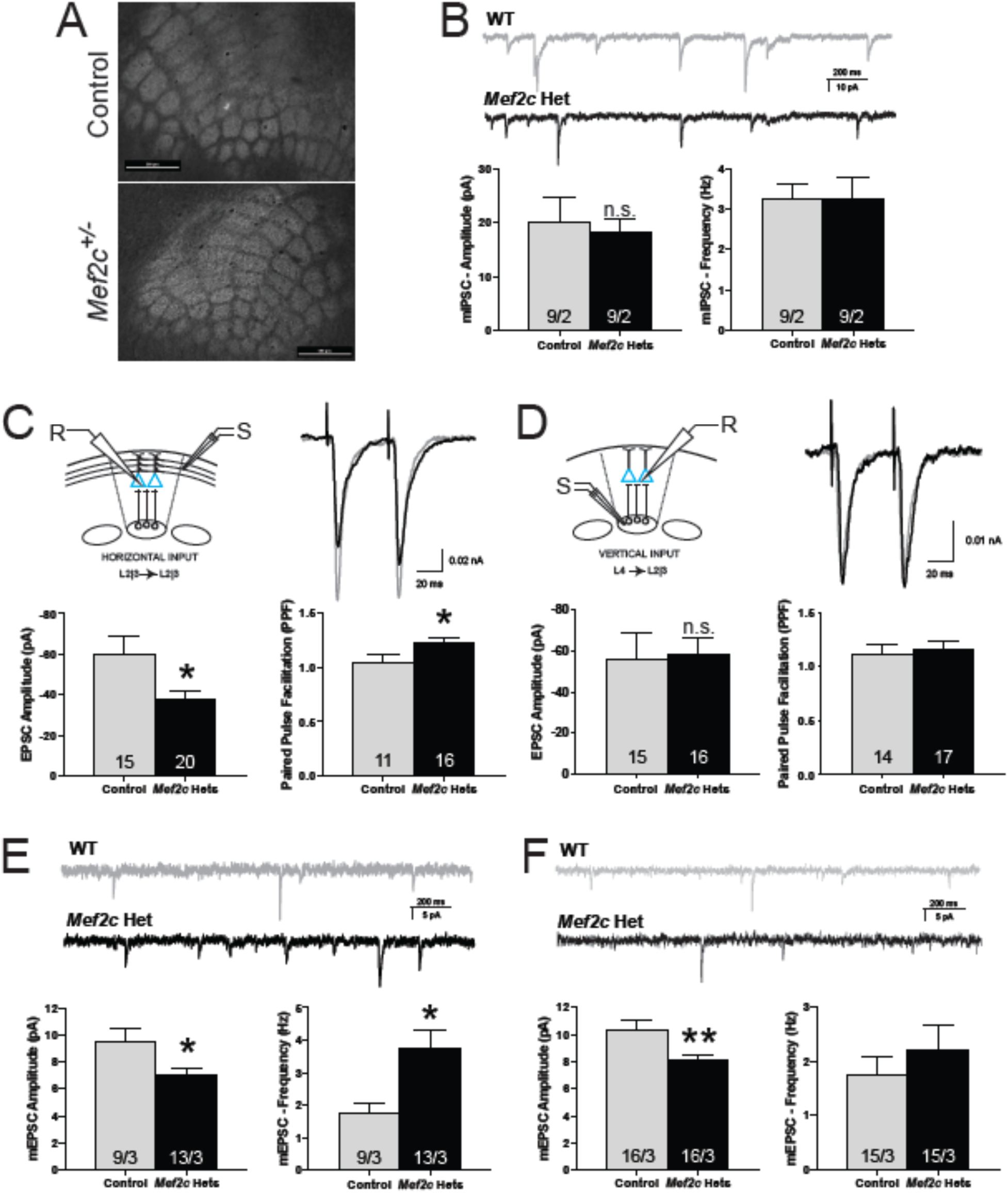
*Mef2c*-Het mice have alterations in cortical synaptic transmission. (A) Both control and *Mef2c*-Het mice have normal barrel fields in the somatosensory cortex, as reflected by VGlut2 staining. Scale bar=500 μm. (B-F) Ex vivo recordings from organotypic slices were collected from pyramidal neurons within the barrel cortex field. (B) No changes were observed in mIPSC amplitude or frequency in the *Mef2c*-Het layer 2/3 pyramidal neurons. (C) Reduced EPSC amplitude and increased paired pulse facilitation (PPF) were observed in layer 2/3 *Mef2c*-Het neurons after stimulating input neurons from neighboring layer 2/3 neurons in adjacent barrel fields (horizontal inputs). (D) No changes in evoked EPSC amplitude or PPF were observed in layer 2/3 pyramidal neurons after stimulating input neurons from layer 4 (vertical inputs). “R” is recording electrode. “S” is stimulating electrode. (E,F) *Mef2c*-Het cortical pyramidal neurons have reduced mEPSC amplitude in layer 2/3 (E) and layer 5 (F), and increased mEPSC frequency in layer 2/3 (E). Data are reported as mean ± SEM. Statistical significance was determined by unpaired t-test. *p<0.05. Number of cells and animals, respectively, are reported in each graph. Also see Figure S3.

### Mef2c-Het mice display dysregulation of cortical genes associated with ASD risk, excitatory neurons and microglia

Using an unbiased RNA-sequencing (RNA-Seq) approach, we examined gene expression from whole cortex in control and *Mef2c*-Hets (p35-p40), and we identified 490 genes that were significantly dysregulated (FDR < 0.05; Fig. 4A, S4A; Table S1,S2). We confirmed the differential expression of a number of interesting *Mef2c*-Het differentially expressed genes (DEGs) that are associated with ASD risk, microglia, and others by qRT-PCR (Fig. 4D). We also investigated the association of *Mef2c*-Het DEGs with genetic and genomic data from various brain disorders. We found that the *Mef2c*-Het DEGs, particularly the downregulated genes, were overrepresented in genes associated with ASD risk and FMRP binding (Fig. 4B; Table S1; Fig. 4D). We also assessed enrichment for *Mef2c*-Het DEGs in genes that are dysregulated in a meta-analysis of transcriptomic data across neuropsychiatric disorders (Gandal et al., 2018). Interestingly, *Mef2c*-Het DEGs, particularly the downregulated genes, were significantly enriched for a PsychENCODE excitatory neuron module of genes that are downregulated in ASD (versus other neuropsychiatric disorders) brains (geneM1; Fig. 4C; Table S2). We also observed significant enrichment of *Mef2c*-Het DEGs in PsychENCODE gene module 8 (geneM8; Fig. 4C; Table S2), which is an excitatory neuron module of genes downregulated in both ASD and schizophrenia and enriched for Bipolar Disorder (BPD) genetic variants. *Mef2c*-Het DEGs, particularly the upregulated genes, were enriched in PsychENCODE module 6, which is a microglia module of genes upregulated in ASD, but downregulated in SCZ and BPD (geneM6; Fig. 4C; Table S2). Using single-cell RNA-seq data from mouse cortex (Saunders et al., 2018), we observed that *Mef2c*-Het DEGs were strongly enriched for cortical excitatory neuron genes and microglia genes (Fig. S4B; Table S2), further supporting the importance of MEF2C in regulating gene expression in the two key brain populations where MEF2C expression is most highly expressed.

**Figure 4.**
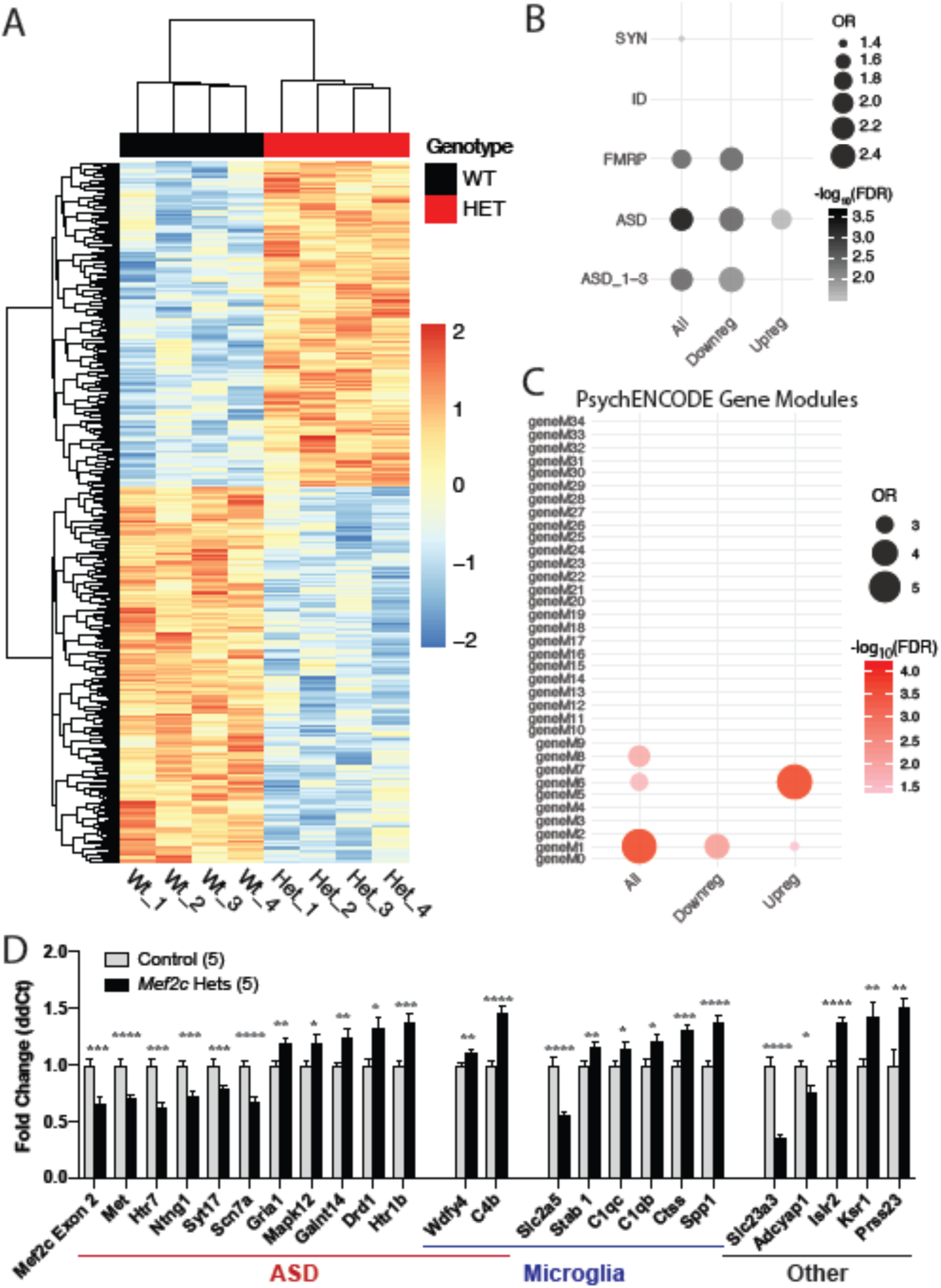
Differentially expressed genes in *Mef2c*-Het cortex. (A) Heatmap showing differentially expressed genes (DEGs) in *Mef2c*-Het cortex (p35-p40) compared with controls. In red, are genes with higher expression; in blue, are genes with lower expression. (B) *Mef2c*-Het DEGs are significantly enriched in genes associated with FMRP, ASD, or scored ASD (ASD_1-3; high-confidence ASD genes) (see Methods). (C) *Mef2c*-DEGs are enriched in gene modules dysregulated in neuropsychiatric disorders, specifically the M1 and M6 modules. (D) qPCR validation of select *Mef2c-*Het DEGs associated with autism, microglia, or other cellular functions. Data are reported as mean ± SEM (D). Statistical significance was determined by unpaired t-test (D). *p<0.05, **p<0.01, ***p<0.005, ****p<0.0005. See Methods for statistical analysis of A-C. Number of animals (n) is 4/genotype for RNA-Seq and 5/genotype for qPCR validation. Also see Figure S4.

Gene ontology analysis of the *Mef2c*-Het DEGs revealed significant enrichment of microglia proliferation genes, cell metabolism genes, and genes in a microglia subpopulation in the developing brain that is restricted to unmyelinated axon tracts (Fig. S5). Since *Mef2c*-Hets showed significant dysregulation of microglial genes (Fig. 4C,D), and MEF2C is expressed in microglia in the developing and mature brain (Fig. S5) (Deczkowska et al., 2017; Gosselin et al., 2017; Zhang et al., 2014a), we analyzed the *Mef2c*-Het brain for possible upregulation of the microglia cell-type and neuroimmune activation marker, ionized calcium-binding adapter molecule 1 (Iba1) (Ito et al., 1998; Ito et al., 2001). In both the cortex and hippocampus, we observed a significant increase in Iba1 expression (Fig. 5A-C,E; ****p<0.0001, K-S test) without a change in the density of microglia (Fig. 5D,F), suggesting possible microglial activation in the *Mef2c*-Het brain. This increase in Iba1 was present without an obvious change in microglial cell morphology or microglial cell soma volume (Figs. 5A,B,G). In addition, in the *Mef2c-*Het cortex, we observed no changes in classic- and alternative-pathway pro-inflammatory genes, including IL-1B, TNF*α*, CD68, IFN*γ*, and several others (Fig. 5G). However, we did note a significant increase in the expression of several complement-related genes linked previously to synaptic pruning and/or ASD risk, including *C1qb, C1qc* and *C4b* (Fig. 4D, (Bialas and Stevens, 2013; Odell et al., 2005; Schafer et al., 2012; Sekar et al., 2016; Stevens et al., 2007). Moreover, we observed significant enrichments of upregulated *Mef2c*-Het DEGs in scRNA-seq gene clusters associated with embryonic-like microglia, postnatal immature microglia, and homeostatic microglia (Fig. 5H). Taken together, these results reveal that the reduction of MEF2C levels has significant impacts on microglia gene expression programs.

**Figure 5.**
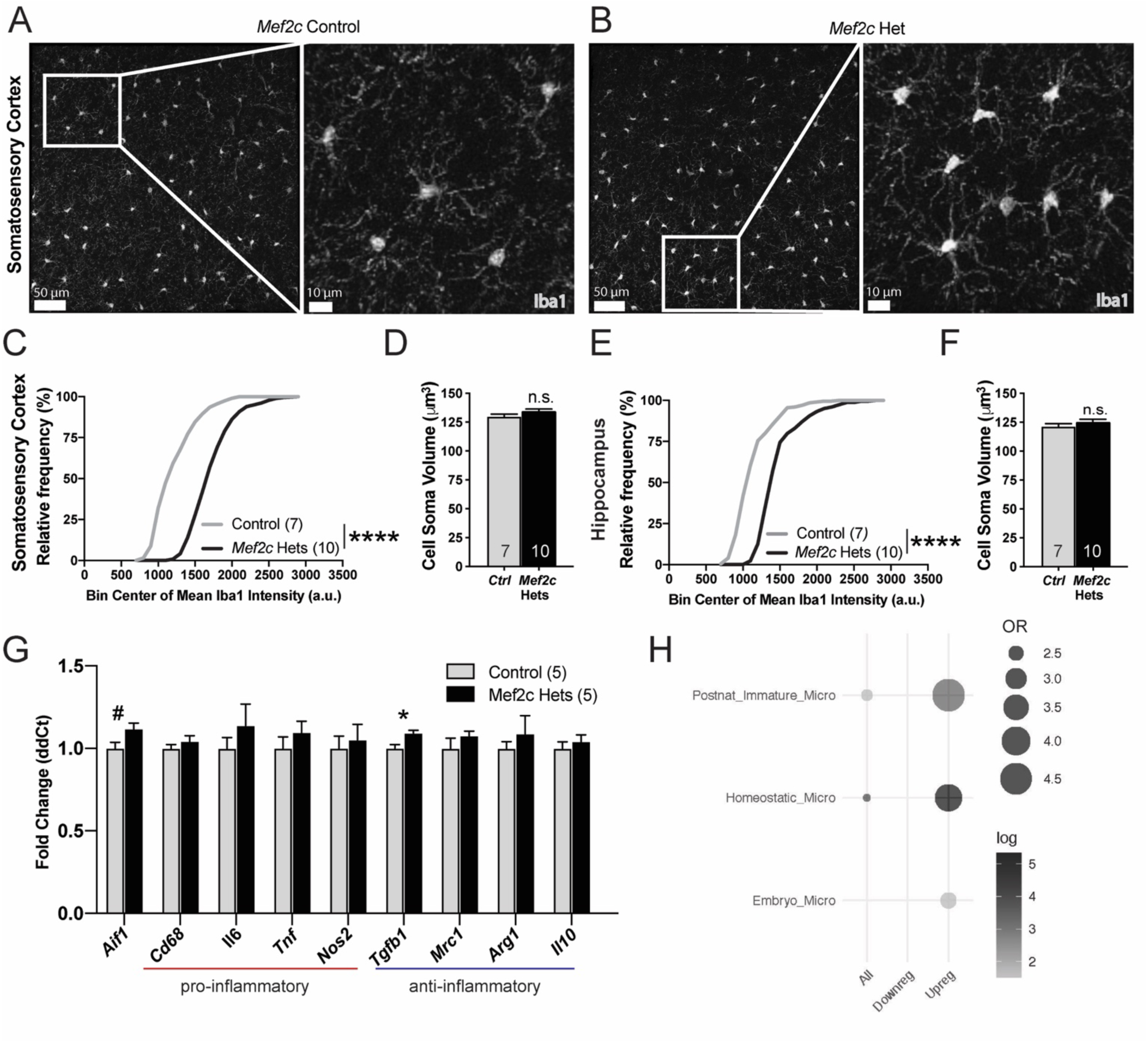
*Mef2c-*Het mice exhibit increased Iba1 expression levels. (A,B) Representative images of Iba1-positive microglia in the SSCtx in control (A) and *Mef2c*-Het mice (B). (C,E) *Mef2c*-Het mice have a right-shifted cumulative frequency distribution of mean Iba1 intensities in Iba1-positive cells (microglia) in the SSCtx (C) and hippocampus (E) compared to controls. Gray line represents distribution of control cells and black line represents distribution of *Mef2c-*Het cells. (D,F) There is no difference in the cell soma volume of Iba1 positive cells (microglia) in the SSCtx (D) or hippocampus (F) between controls and *Mef2c-*Het mice. (G) Fold changes of genes associated with microglial activation in controls and *Mef2c*-Hets. (H) Mef2c-Hets have an upregulation of genes expressed in postnatal immature, homeostatic, and embryonic microglia. Unless specified, data are reported as mean ± SEM. Statistical significance determined by Kolmogorov–Smirnov test (C,E) or unpaired two-tailed nested t-test (D,F), unpaired two-tailed t-test (G). ****<0.0001. Sample sizes for each genotype are denoted on bars of or above each graph unless otherwise specified. Images (A,B) have contrast and brightness enhanced for ease of viewing. Images are modified equally for both genotypes. Also see Figure S5.

### MEF2C contributes to neurotypical behaviors through key roles in forebrain excitatory neurons and microglia

In the developing and mature mouse brain, MEF2C is expressed in several neuronal cell types, including cortical excitatory pyramidal cells and Parvalbumin-positive GABAergic inhibitory neurons, and in microglia (Barbosa et al., 2008; Harrington et al., 2016; Kamath and Chen, 2018; Mayer et al., 2018; Zhang et al., 2014b). Since the *Mef2c*-Het mouse cortex showed robust changes in both excitatory neurons and microglia gene expression (Figs. 4, S4), we generated cell type-specific conditional *Mef2c* heterozygous mice to explore the contribution of forebrain excitatory neurons versus microglia for the development of MCHS-like phenotypes. We first generated mice heterozygous for *Mef2c* in Emx1-lineage cells (*Mef2c*-cHet*^Emx1-cre^*) (Iwasato et al., 2008), which represents ∼85% of forebrain excitatory neurons throughout the cortex and hippocampus. Similar to the global *Mef2c*-Hets, the *Mef2c*-cHet*^Emx1-cre^*mice displayed altered anxiety-like behavior and male-selective increases in locomotion and repetitive jumping (Fig. 6A-C), but they showed no changes in social behavior or shock sensitivity (Fig. 6D, S6A). Interestingly, similar to global *Mef2c*-Hets (Fig. 3E) and *Mef2c* cKO*^Emx1-cre^* mice (Harrington et al., 2016), we observed a reduction of mEPSC amplitude in layer 2/3 pyramidal neurons from *Mef2c*-cHet*^Emx1-cre^*mice (Fig. S6D), suggesting that the deficit in cortical glutamatergic synaptic transmission in layer 2/3 of the global *Mef2c*-Hets is due to MEF2C’s function in excitatory forebrain neurons. In addition, mice heterozygous for *Mef2c* in PV-positive GABAergic interneurons (*Mef2c*-cHet*^PV-cre^*) – a population of predominantly GABAergic interneurons with high MEF2C expression – also showed male-selective increases in locomotion (Fig. 6F), but these mice did not develop changes in repetitive jumping, anxiety-like behavior, social interaction, or shock sensitivity (Figs. 6E,G-H, S6B). These findings suggest that Emx1-lineage excitatory forebrain neurons, and to a much lesser extent PV-expressing neurons, contribute to the development of some, but not all, of the behavior phenotypes observed in the global *Mef2c*-Hets.

**Figure 6.**
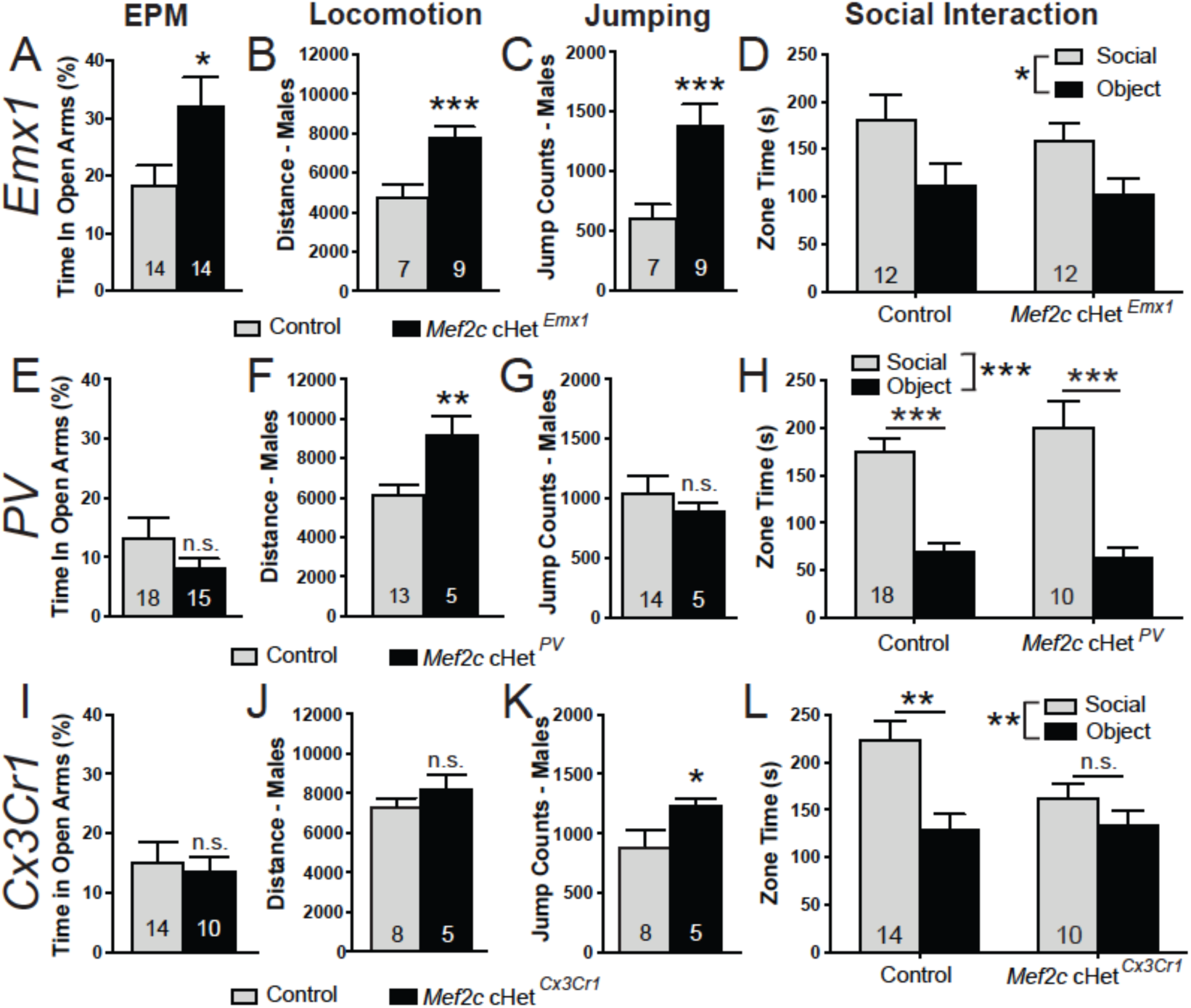
Cell type-selective behavior phenotypes in *Mef2c* conditional heterozygous (*Mef2c*-cHet) mice. (A-D) Behaviors in *Mef2c* cHet*^Emx1^*mice. (A) *Mef2c* cHet*^Emx1^* mice spend more time on the open arms of the elevated plus maze. (B,C) Male *Mef2c*-cHet *^Emx1^* mice are hyperactive (B) and show increased jump counts (C). (D) *Mef2c* cHet*^Emx1^* mice have normal social interaction. (E-H) Behaviors in *Mef2c* cHet*^PV^*mice. (E) *Mef2c* cHet*^Pcp2^* mice spend similar time on the open arms of the EPM. (F,G) Male *Mef2c* cHet*^PV^*mice are hyperactive (F) but show normal jump counts (G). (H) *Mef2c* cHet*^PV^* mice have normal social interaction. (I-L) Behaviors in *Mef2c* cHet*^Cx3cr1^* mice. (I) *Mef2c* cHet*^Cx3cr1^* mice are similar to controls in elevated plus maze. (J,K) Male *Mef2c* cHet*^Cx3cr1^* mice have normal activity (J) but show increased jump counts (K) compared to control mice. (L) *Mef2c* cHet*^Cx3cr1^* mice have a lack of preference for interacting with a novel mouse (social) over the novel object. Data are reported as mean ± SEM. Statistical significance was determined by unpaired t-test (A-C,E-G,I-K) or 2-way ANOVA (D,H,L). n.s.=not significant, *p<0.05, **p<0.01, ***p<0.005, n.s. = not significant. Number of animals are reported in each graph. Also see Figure S6.

To explore a possible contribution of MEF2C in microglia to MCHS-like behaviors, we generated mice heterozygous for *Mef2c* selectively in microglia (*Mef2c*-cHet*^Cx3cr1-cre^*). The conditional mutant mice displayed social impairments in the 3-chamber social interaction test (Fig. 6L, *p<0.05, two-way ANOVA), similar to the *Mef2c*-Het mice. In addition, *Mef2c*-cHet*^Cx3cr1-cre^* mice showed a significant increase in male-specific repetitive jumping (Fig. 6K, *p<0.05, Welch’s t-test), but with no discernable effects on exploratory activity (Fig. 6J), anxiety-like behavior or shock sensitivity (Fig. 6I, S6C). Taken together, our observations suggest that *Mef2c* haploinsufficiency in early postnatal microglial cells is sufficient to produce autism-related behaviors, and that the majority of MCHS-like phenotypes in the global *Mef2c*-Hets are a consequence of MEF2C hypofunction in both neurons and microglia.

## Discussion

We report here three new *MEF2C* mutations identified in individuals with MCHS-related symptoms, and these *MEF2C* missense or small duplication mutations disrupted MEF2C DNA binding. Interestingly, all of the known MCHS missense or duplication mutations cluster within the highly-conserved MADS and MEF2 domains (Fig. 1A) that mediate the DNA binding and dimerization functions of MEF2C (McKinsey et al., 2002). We also found that global, DNA binding-deficient *Mef2c* heterozygous mice display numerous behavioral phenotypes reminiscent of MCHS, including social interaction deficits, ultrasonic vocalization deficits, motor hyperactivity, repetitive behavior, anxiety-related behavior and reduced sensitivity to a painful stimulus (footshock). The *Mef2c-*Hets also possessed input-selective pre- and postsynaptic deficits in glutamatergic excitatory synaptic transmission in the somatosensory cortex. Gene expression analysis of cortical tissue from *Mef2c*-Hets revealed significant enrichment of genes linked to ASD risk, excitatory neurons and microglia, which is notable considering the enrichment of dysregulated genes linked to cortical excitatory neurons and microglia in brains of individuals with idiopathic ASD (Velmeshev et al., 2019). Conditional *Mef2c* heterozygous mice in Emx1-lineage cells, which represent predominantly forebrain excitatory neurons, reproduced several of the global *Mef2c*-Het behaviors and cortical synaptic phenotypes. Consistent with the dysregulation of microglial genes in *Mef2c*-Het mice, early postnatal conditional *Mef2c* heterozygosity in *Cx3cr1*-lineage cells, which in the brain are almost exclusively microglia (Hoogland et al., 2015; Ito et al., 1998; Ito et al., 2001), produced offspring with social deficits and increased repetitive behavior. Our findings support the emerging view that microglial dysfunction in the developing brain can contribute to the development of ASD symptoms.

While our findings reveal a key role for MEF2C in both neurons and microglia for neurotypical behaviors in mice, there are a number of new questions raised by these findings. For example, why do we see male-selective effects of *Mef2c* heterozygosity on hyperactivity and/or jumping behavior in the *Mef2c*-Het and *Mef2c*-cHet mice (Figs. 2E,F; 6B,C,F,K)? This suggests an interaction between sex-based mechanisms of development and MEF2C-dependent transcription during development, and indeed, numerous studies have demonstrated that both neuron and microglia functions can be differentially regulated in males and females (Wright-Jin and Gutmann, 2019). It is also interesting to note that *Mef2c-*Het DEGs linked to excitatory neurons show a preferential downregulation, whereas *Mef2c*-Het DEGs linked to microglia display a preferential upregulation. MEF2C is reported to function as both a transcriptional activator and a repressor, and there are cell type-specific signaling mechanisms that regulate MEF2C activity(Harrington et al., 2016; Kang et al., 2006; Lyons et al., 2012). Determining whether or how MEF2C regulates cell type-specific gene expression as an activator or repressor, or whether the DEGs are an indirect consequence of MEF2C hypofunction, will be important goals for future studies.

MEF2 proteins can regulate activity-dependent glutamatergic synapse elimination (Flavell et al., 2006; Pfeiffer et al., 2010; Pulipparacharuvil et al., 2008; Tsai et al., 2012). MEF2C can function in cortical pyramidal neurons as a cell-autonomous transcriptional repressor to regulate dendritic spine density, synapse number and AMPA-mediated postsynaptic strength (Harrington et al., 2016; Rajkovich et al., 2017). Conditional deletion of both *Mef2c* gene copies in forebrain excitatory neurons produces mice with dramatic changes in cortical synapse functions, including decreased glutamatergic synaptic transmission, and numerous alterations in typical mouse behaviors, including learning and memory, reward-related behavior, motor hyperactivity and repetitive behaviors, and differential gene expression (Adachi et al., 2015; Barbosa et al., 2008; Harrington et al., 2016; Li et al., 2008). In contrast to these *Mef2c* brain conditional knockouts, humans with MCHS possess deletions or mutations in a single gene copy throughout the entire body (Berland and Houge, 2010; Bienvenu et al., 2013; Engels et al., 2009; Le Meur et al., 2010; Mikhail et al., 2011; Novara et al., 2010; Paciorkowski et al., 2013; Tonk et al., 2011; Vrecar et al., 2017; Zweier et al., 2010; Zweier and Rauch, 2012). Since MCHS symptoms are reported predominantly from macro- and microdeletions that disrupt *MEF2C* and other neighboring genes, we sought to identify possible loss-of-function *MEF2C* mutations within its protein coding region to better understand the relationship between symptoms and *MEF2C*. By comparing multiple new *MEF2C*-related mutations from individuals with developmental delay and other MCHS-associated symptoms, we observed that all of the missense mutations resided within the MEF2C DNA binding and dimerization domains (MADS/MEF2). In addition, there were multiple mutations that produced a premature stop codon or a frameshift predicted to produce a truncated MEF2C lacking the C-terminal nuclear localization sequence. One interesting observation was the ∼1.3-fold increase in *Mef2c* mRNA levels in the *Mef2c*-Het mouse cortex (Fig. 4). However, there was a ∼50% reduction of the exon 2-containing transcripts (the Cre-flox edited exon), resulting in ∼35% overall reduction in full-length, functional MEF2C (Fig. 4D). This suggests that MEF2C might negatively regulate its own gene expression, as originally reported in muscle cells (Wang DZ et al., 2001), and that a relatively modest overall reduction in MEF2C levels is sufficient to produce numerous changes in gene expression, synaptic development and function, and neurotypical behaviors.

Developing and mature microglia play important roles in brain development, including synaptic phagocytosis (Paolicelli et al., 2011; Schafer et al., 2012). Microglia also mediate synapse patterning, neurogenesis, myelinogenesis, and cellular phagocytosis (Parkhurst et al., 2013; Sierra et al., 2010; Zhan et al., 2014). MEF2C is expressed in both human and mouse microglia, and MEF2 proteins regulate microglia development (Gosselin et al., 2017). Microglia-enriched RNAs are dysregulated in human cortex from idiopathic ASD brains (Velmeshev et al., 2019) and in the mouse *Mef2c*-Het cortex (Fig. 4, S4), and we find that *Mef2c* hypofunction in microglia (Cx3cr1-lineage) is sufficient to produce autism-like behaviors in mice (Fig. 6K,L). Interestingly, despite a strong increase in the *Mef2c*-Het brain of the microglia cell-type and activation marker, Iba1, (Fig. 5) as well as other microglia genes including several complement genes (*e.g. C1qb*, *C1qc* and *C4b*), osteopontin (*Spp1*), and Cathepsin S (*Ctss*) (Fig. 4C,D), we failed to detect a clear signature of neuroinflammation (Fig. 5G). Our findings suggest that loss of one *Mef2c* copy does not produce classic microglial “activation”, but rather that microglial development, function or maturation might be perturbed. Of note, *Mef2c*-Het DEGs showed enrichment for a scRNA-seq cluster of genes associated with embryonic and immature postnatal microglia, suggesting a possible delay in microglia maturation in the *Mef2c*-Het mice. Future studies will be important to determine the precise roles of MEF2C in microglial development and function, and whether *Mef2c* heterozygosity alters one or more of the numerous reported roles for microglia in brain development.

Taken together our findings reveal that MEF2C hypofunction throughout development produces numerous complex changes in cortical synaptic transmission, gene expression and behaviors reminiscent of MCHS and ASD. Specifically, the *Mef2c*-Het behaviors are associated with robust, input-selective deficits in cortical excitatory synaptic transmission, and disruption of excitatory neuronal and microglial gene expression. Importantly, our cell type-selective manipulations strongly suggest that MEF2C contributes to neurotypical development through critical roles in multiple cell types, including forebrain excitatory neurons (Emx1-lineage), PV-positive interneurons and microglia. Understanding the role of MEF2C in these, and other, cell populations in the body are likely to provide important new insights into effective treatment strategies for symptoms of MCHS.

## Acknowledgments

This work was supported in part by NIH grant R01 MH111464 (to C.W.C.), TL1 TR001451, UL1 TR001450 and F30 HD098893 (to C.M.B.), the Brain and Behavior Research Foundation NARSAD Young Investigator Award (to A.J.H.) and Simons Foundation SFARI pilot grant #649452 (to C.W.C.). The authors would like to acknowledge Dr. Patrick J. Mulholland and the Shared Confocal Core (NIH grant S10 OD021532), Dr. Jeremy L. Barth and the MUSC Proteogenomics Facility for qPCR instrument (supported by NIGMS GM103499 and MUSC’s Office of the Vice President for Research), and Duncan Nowling for technical assistance.

## Author Contributions

A.J.H., C.M.B., A.A., and C.W.C. designed experiments, performed data analysis, and wrote the manuscript. H.W.M, D.B.E., and S.A.S. collected MCHS patient data. A.J.H., C.M.B., A.A., K.B., and Y.J.C. performed behavior test and analyzed data. S.B. and G.K. analyzed RNA-Seq data. E.T. performed electrophysiology and data analysis. B.M.S. and M.D.S. performed dendritic spine morphology experiments. A.J.H., C.M.B., K.B., Y.J.C., and A.T. performed molecular/biochemical experiments and data analysis. A.J.H. and C.M.B. performed statistical analyses.

## Supplemental Information

### Supplemental Figures

**Supplemental Table S1. RNA-Seq gene expression in control and *Mef2c*-Het cortex. See file “Mef2c het Database_GenSet.”**

**Supplemental Table S2. *Mef2c* DEGs compared to single-cell RNA-Seq gene databases. See file “Mef2c het Database_single-cell.”**

**Supplemental Table S3.**
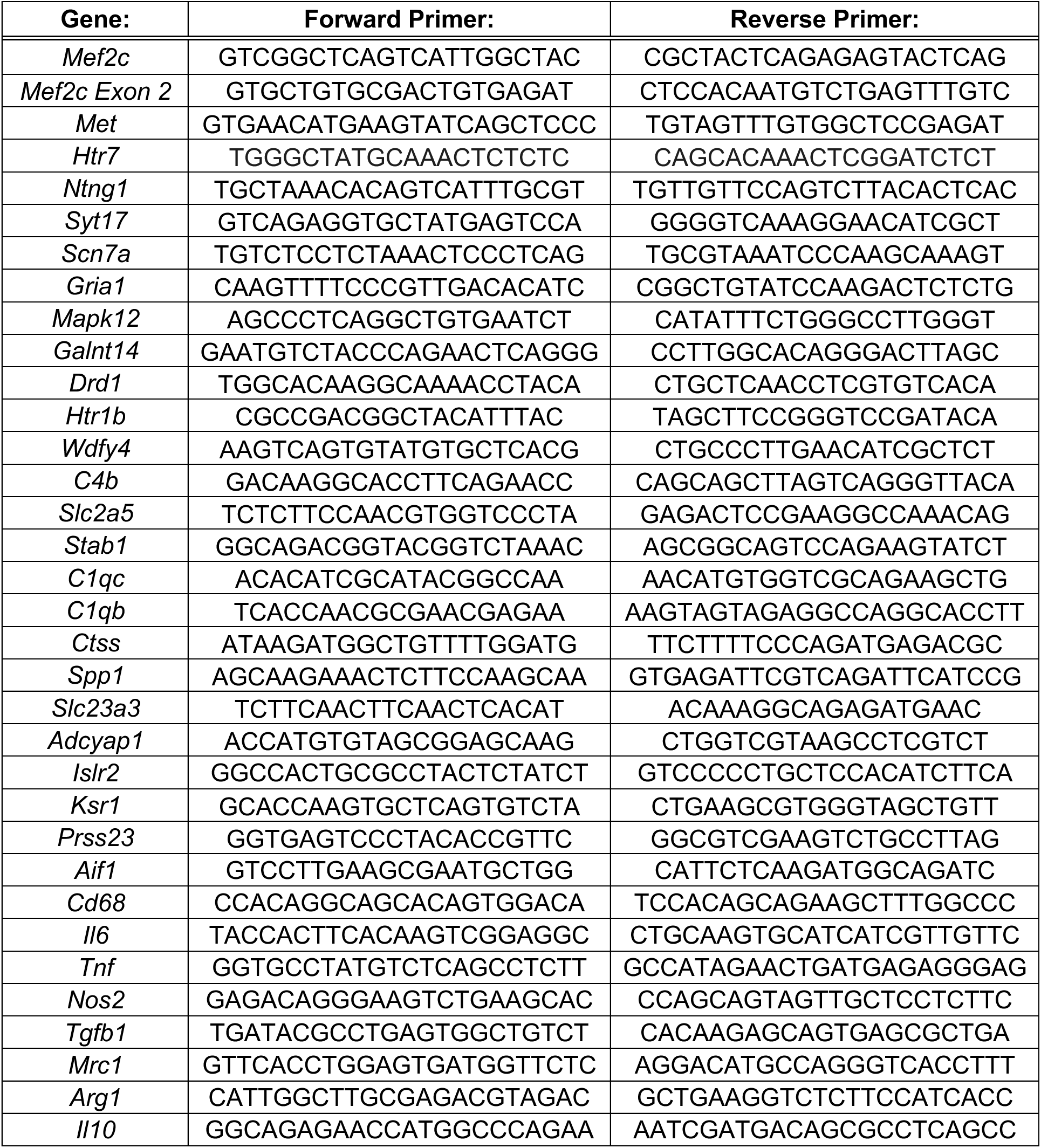
Primers for qPCR analysis of *Mef2c*-Het DEGs.

**Figure S1.**
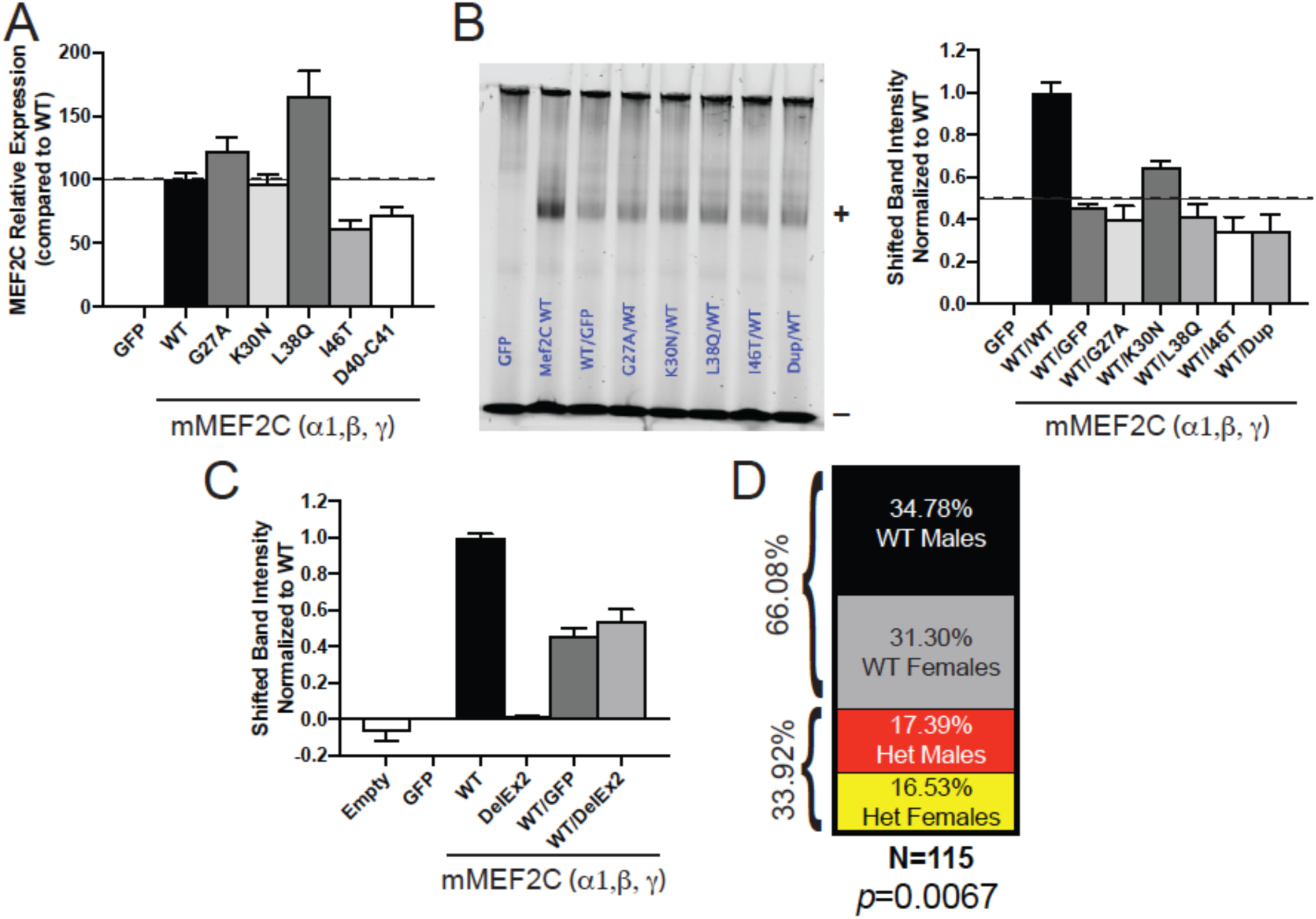
(A) Quantification of MEF2C Western blot from 293-T cells containing MCHS mutations (Fig. 1B); n=3. (B) EMSA shows that MCHS-mutations in MEF2C do not disrupt WT MEF2C from binding to the MEF2 response element (MRE); n=5. Quantification of MEF2C bound probe is reported (B). (C) Quantification of MEF2C DelEx2 EMSA (Fig. 1F); n=4. (D) Genetic distribution of offspring from WT and *Mef2c*-Het mice (n=115). Data are reported as mean ± SEM. Statistical significance was determined by Chi-squared test (D). Also see Figure 1.

**Figure S2.**
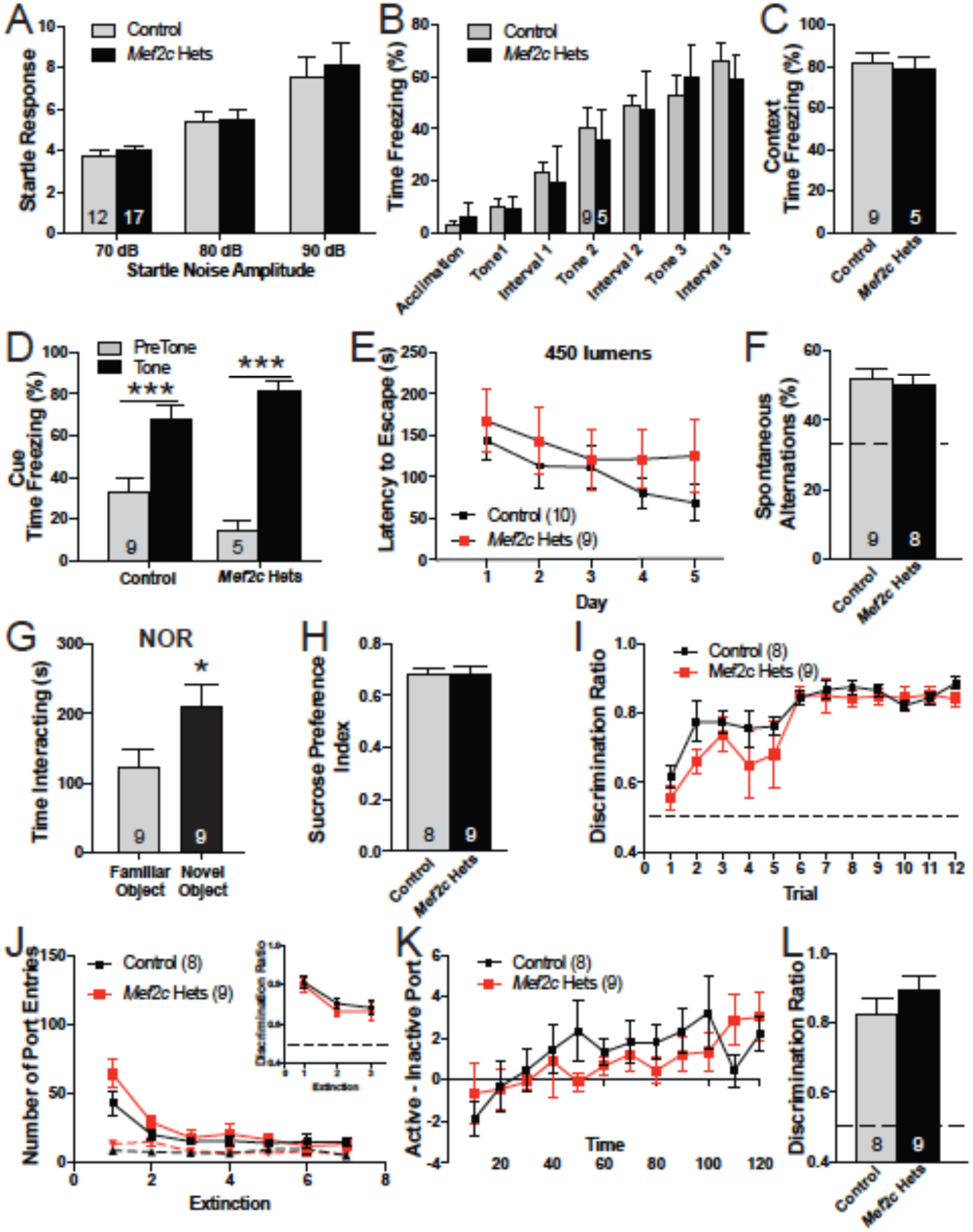
*Mef2c*-Het mice have normal cognitive abilities. (A) There is no difference in acoustic startle response between *Mef2c*-Het and control mice. (B-D) Pavlovian Fear Conditioning. Both control and *Mef2c*-Het mice increase freezing with each tone/shock pairing during training (B) and show similar levels of freezing during the context (C) and cue (D) test. (E) No difference in latency to escape a very bright (450 lumens) Barnes Maze. (F) Both control and *Mef2c*-Het mice have similar spontaneous alternations in the Y-maze. (G) Novel object recognition. *Mef2c*-Het mice (n=9) interacted more with a novel object than a familiar object. (H) Both genotypes show a similar preference for 1% sucrose solution. (I-L) Sucrose Self-Administration Task. (I) Discrimination ratio during sucrose self-administration. (J) Number of active (solid line) and inactive (dashed line) port entries during extinction. Control and *Mef2c*-Het mice can recall the active port on the first day of extinction, and both genotypes show similar extinction rates (J) and discrimination ratio (insert). (K) Discrimination ratio during the first day of sucrose self-administration. (L) Both control and *Mef2c*-Het mice have high discrimination ratios during cue-induced reinstatement of sucrose seeking. Data are reported as mean ± SEM. Statistical significance was determined by 2-way ANOVA (A,B,D,E,I-K) or unpaired t-test (C,F-H,L). *p<0.05, ***p<0.005. Number of animals (n) are reported in each graph for respective experiment. Also see Figure 2.

**Figure S3.**
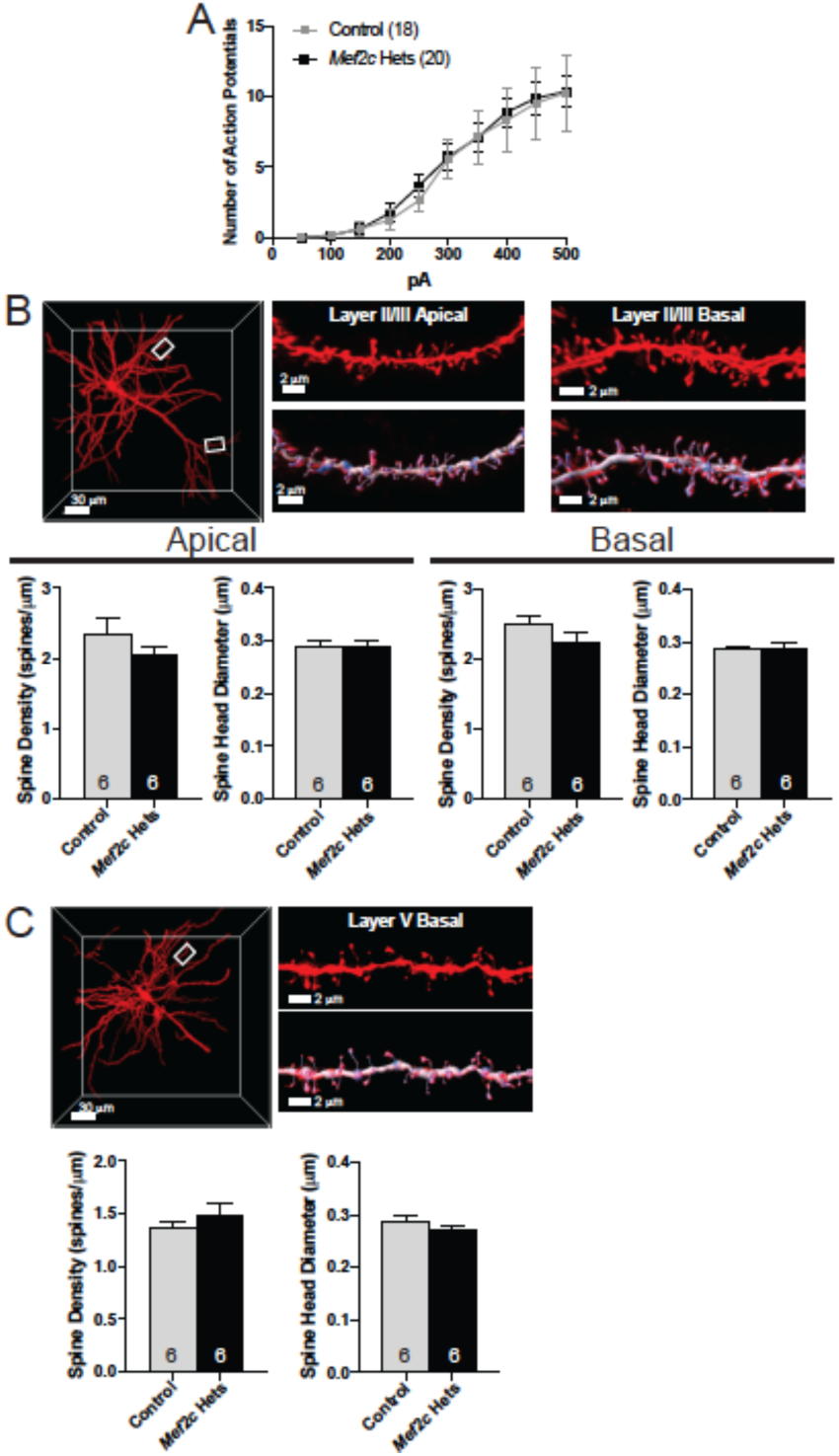
(A) Both control and *Mef2c*-Het cortical pyramidal neurons have similar numbers of action potentials evoked by increasing current injections (500ms, 50pA steps) recorded in current clamp. (B,C) Representative image of a DiI filled cortical pyramidal neuron (B=layer 2/3; C=layer 5) and representative apical and basal dendritic segments used for quantifying dendritic spines. Both control and *Mef2c*-Het mice have similar dendritic spine density and spine head diameter on apical and basal dendrites. Data are reported as mean ± SEM. Statistical significance was determined by 2-way ANOVA (A) or unpaired t-test (B,C). Number of cells (A) or animals (B,C) are reported in each graph for respective experiment. Scale bar=30 μm (neuron) or 2 μm (dendritic stretch). Also see Figure 3.

**Figure S4.**
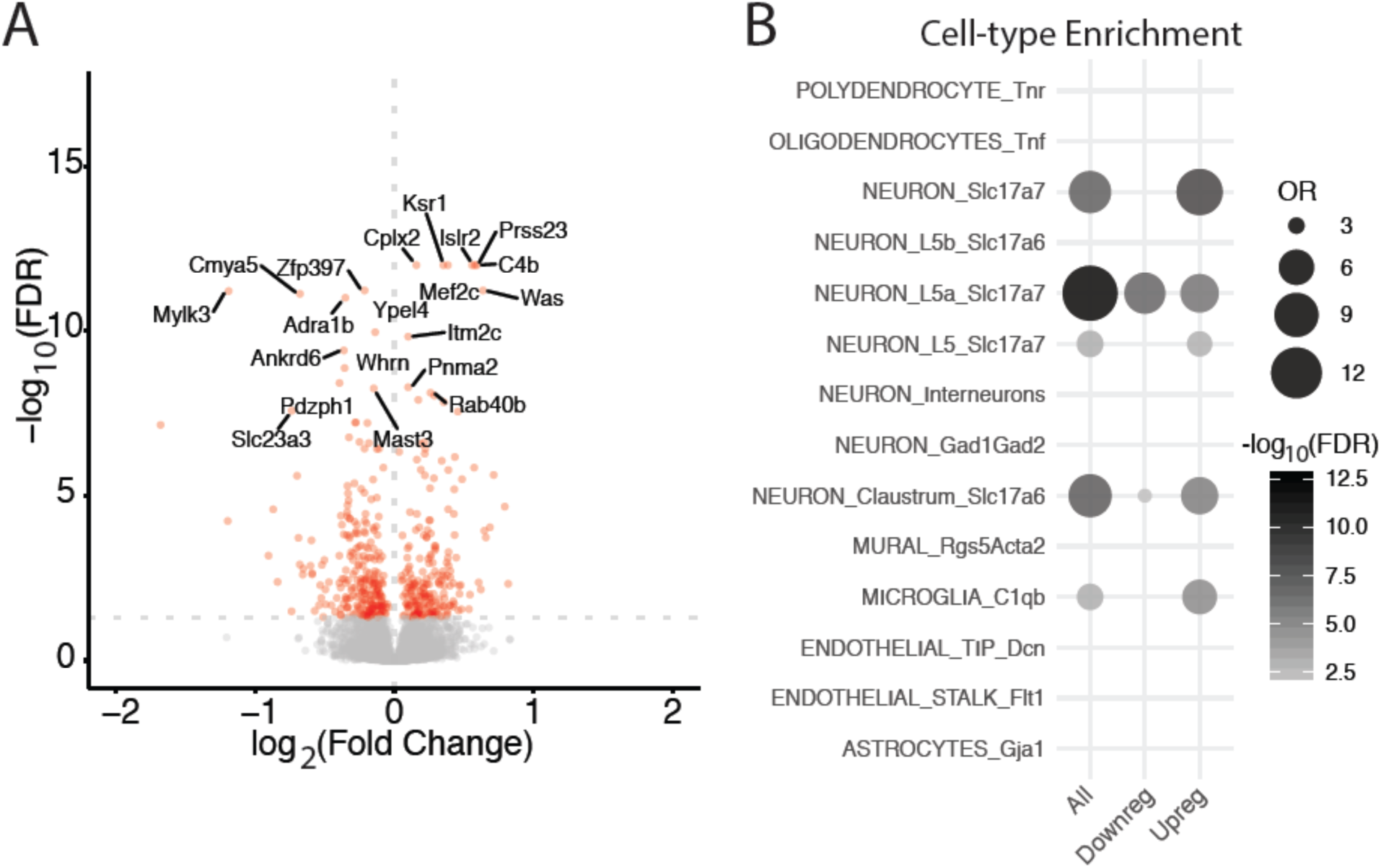
Differentially expressed genes in *Mef2c*-Het cortex. (A) Volcano plot of *Mef2c*-Het DEGs shows genes are both up- and down-regulated in the cortex of *Mef2c*-Het mice. (B) Bubble-plot of *Mef2c*-Het DEGs are enriched for genes expressed in neurons and microglia. See Methods for statistical tests. Number of animals is 4/genotype. Also see Figure 4.

**Figure S5.**
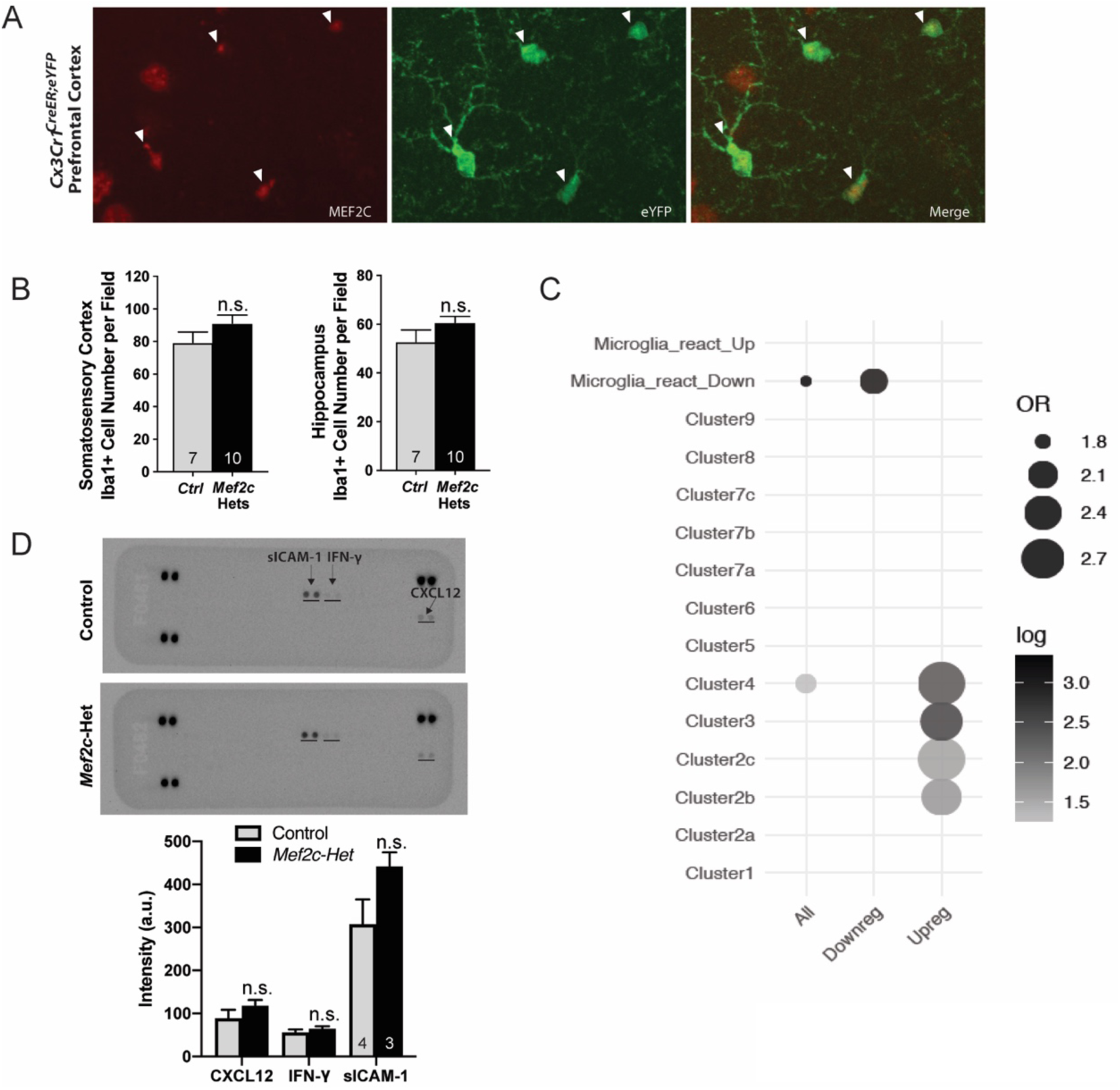
(A) MEF2C is expressed in microglia. Using a microglia reporter mouse (*Cx3Cr1^CreER;eYFP^*), immunohistochemistry reveals that MEF2C (red) is expressed in microglia (green) under basal conditions. (B) Microglial cell density in the somatosensory cortex and dentate gyrus of the hippocampus, quantified via Iba1 staining of microglia, is not significantly different between *Mef2c*-Hets and controls. (C) Enrichment of differentially expressed genes in *Mef2c*-Hets are upregulated in cluster 2b (G-phase proliferative cluster), cluster 2c (M-phase proliferative cluster), and cluster 3 (metabolically active microglia at E14.5), and cluster 4 (microglia associated with unmyelinated axon tracts at (P4/5). (D) Cytokine antibody array analysis for assessing neuroinflammation in *Mef2c*-Hets and controls at P35-P40. Data are reported as mean ± SEM. Statistical significance was determined by unpaired t-test (B,D). Number of animals (B,D) are reported in each graph for respective experiment. Also see Figure 5.

**Figure S6.**
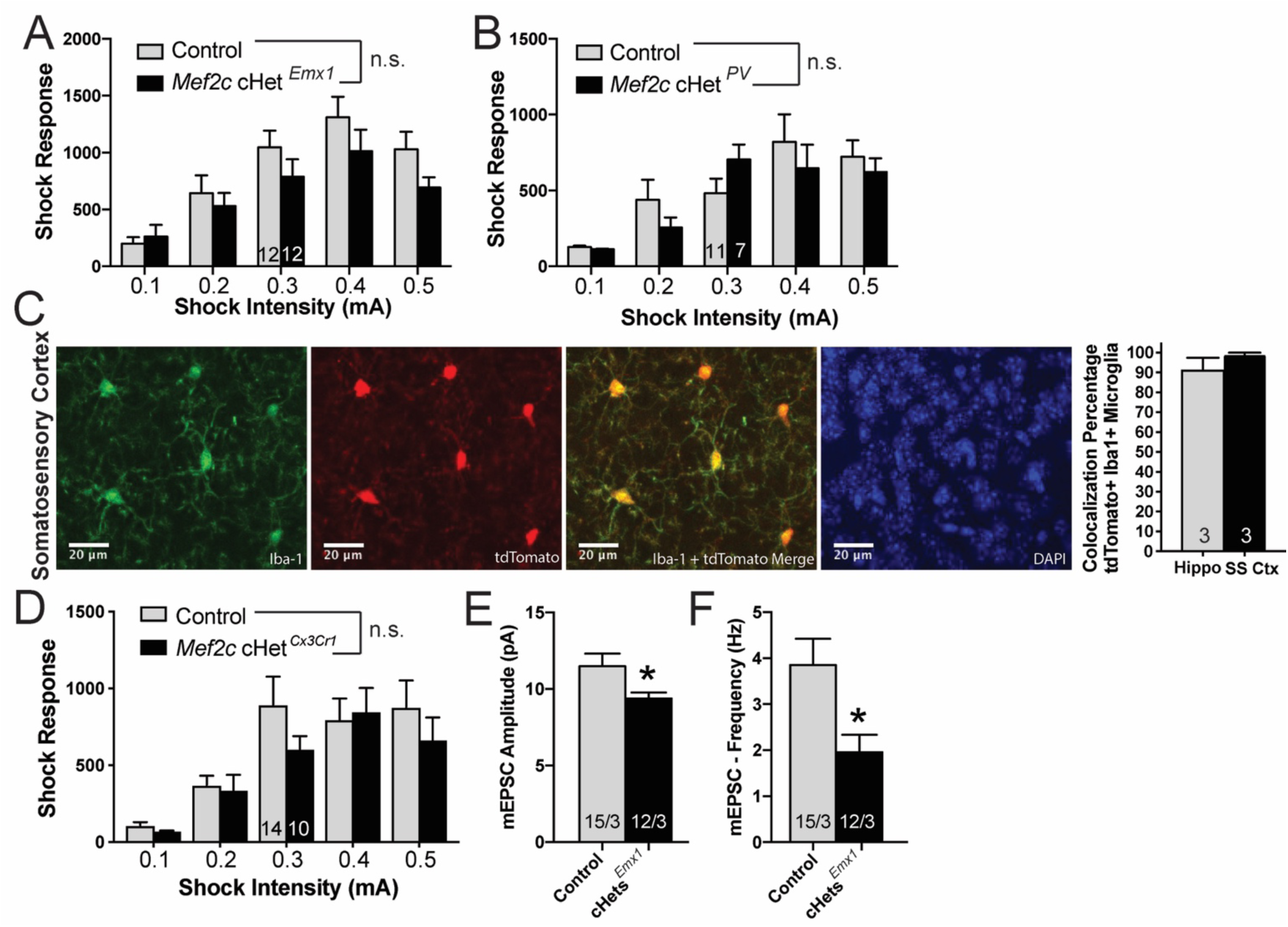
(A,B) *Mef2c* cHet*^Emx1^* (A) and *Mef2c* cHet*^PV^*(B) mice have normal response to shock. (C) Recombination efficiency after p1-p3 tamoxifen treatment in *Cx3Cr1^CreER;eYFP^*; Ai14 mice. tdTomato expression suggests recombination in these Iba1-postive microglia cells. (D) *Mef2c* cHet*^Cx3Cr1^*mice are similar to controls in shock response. (E,F) *Ex vivo* recordings from organotypic slices were collected from pyramidal neurons within the barrel cortex field of *Mef2c* cHet*^Emx1^*mice. Neurons from *Mef2c* cHet*^Emx1^* show reduced mEPSC amplitude (D) and frequency (E) compared to controls. Data are reported as mean ± SEM. Statistical significance was determined by 2-way ANOVA (A-B,D) or unpaired t-test (E,F). Number of animals (A-B,D) or cells/animals (E,F), respectively, are reported in each graph for respective experiment. Also see Figure 6.

## Supplemental Methods

### Animals

Mice (*Mus musculus*) were group housed (2-5 mice/cage; unless specified) with same-sex littermates with access to food and water ab libitum on a 12 hour reverse light-dark cycle. *Mef2c^+/-^* (*Mef2c*-Het) mice were initially generated by crossing *Mef2c^fl/fl^* (RRID:MGI:3719006)(Arnold et al., 2007) mice with *Prm1-Cre* (Jackson Laboratory #003328) to induce germline recombination of *Mef2c*. *Mef2c^+/-^; Prm1-Cre* were crossed with C57BL/6J to remove *Prm1-Cre*, and test mice were generated from *Mef2c^+/-^* mice crossed with C57BL/6J mice. *Mef2c* conditional het mice were generated by crossing *Mef2c^fl/fl^*mice with heterozygous cell-type specific Cre mice (*Emx1-Cre(Iwasato et al., 2008)*; *Pcp2-Cre* (Jackson Laboratory #004146)(Barski et al., 2000); *PV-Cre* (Jackson Laboratory #017320) to generate *Mef2c^fl/+^*; *Cre* conditional heterozygous (*Mef2c* cHet) mice that were compared to their Cre-negative littermates. *Cx3Cr1*-Cre *Mef2c* conditional heterozygous (*Mef2c* cHet^Cx3Cr1^) mice were generated by crossing *Cx3Cr1^creER/creER^* males (Jackson Laboratory #021160)(Parkhurst et al., 2013) with *Mef2c^fl/+^* females to produce *Cx3Cr1^creER/+^*; *Mef2c^fl/+^*(experimental) and *Cx3Cr1^creER/+^*; *Mef2c^+/+^* (control) mice. Experimenters were blinded to the mouse genotype during data acquisition and analysis. Experiments were independently replicated, and the total number of animals/cells were reported in the representative figures. All procedures were conducted in accordance with the Institutional Animal Care and Use Committee (IACUC) and NIH guidelines.

#### EMSA

Electrophoretic mobility shift assay (EMSA) was performed as previously described (Pulipparacharuvil et al., 2008) with modifications. Briefly, proteins were isolated from 293-T cells 48 hours after transfection of pA1T7*α*::*Mef2c* variants in a non-denaturing EMSA cell lysis buffer (20 mM tris-HCl (pH 8.0), 100 mM NaCl, 1 mM EDTA, 1 mM Na_3_VO_4_, 10 mM NaF, 0.5% Nonidet P-40, 1 μM cyclosporin A, 1X Complete Protease Inhibitor cocktail (Roche)), and quantified using the DC protein assay kit (BioRad). Isolated proteins (10 μg) were incubated with fluorescent-tagged (Infrared Dye 700) Mef2 Response Element (IR700::MRE) probes for 60 minutes at room temperature (RT) and resolved in a non-denaturizing acrylamide gel (50 mM tris-HCl (pH 7.5), 380 mM glycine, 2 mM EDTA, 4% acrylamide/bis-acrylamide (BioRad)). Gels were imaged using the Li-Cor Odyssey CLx, and fluorescence was quantified by Li-Cor Image Studio v3.1.4. Data reported represent the mean from at least 3 biological replicates.

#### Immunoblotting

EMSA proteins were denatured by adding 4X sample buffer (+DTT, +*β*ME) to samples in EMSA cell lysis buffer and boiling for 10 minutes. Somatosensory cortex was isolated from adult (8 weeks old) male mice and frozen on dry ice. Tissues were sonicated on ice in an SDS lysis buffer (1% (w/v) SDS, 300 mM sucrose, 10 mM NaF (Sigma), 50 mM HEPES (Sigma), and 1X Complete Protease Inhibitor cocktail (Roche)), boiled for 10 minutes, then centrifuges at 16,000 x g for 10 minutes. Total protein concentration was determined by the DC protein assay kit (BioRad). For western blots, 20 μg of total protein was resolved using 10% SDS-PAGE (BioRad). Proteins were transferred to Immobilon-FL PVDF (Millipore), blocked in Odyssey blocking buffer (Li-Cor) for 2 hours, and incubated overnight with either anti-HA (Sigma-Aldrich H6908; 1:2000) and anti-GFP (Aves Labs GFP-1020; 1:2000) for EMSA proteins, or anti-Mef2c (AbCam ab197070; 1:2500) and anti-Neuronal class III *β*-tubulin (Tuj1, Covance; 1:10,000) antibodies for cortical proteins. Blots were developed with Odyssey CLx Western blot system (Li-Cor Biosciences) (HA, MEF2C, and TUJ1) or ChemiDoc MP Imaging System (BioRad) (GFP).

#### Data Acquisition

All experiments were independently replicated at least twice (typically 3-4 times). The numbers of animals/neurons/dendritic stretches are reported in each figure, and these numbers were estimated based on previous reports. Outliers were determined using GraphPad’s Outlier calculator (Grubb’s) and excluded from data analysis.

#### Mouse Behavior Testing

For all behavior tests, test mice were acclimated to transport and handling for at least 3 days prior to testing. Before each test, mice were acclimated after transport for >30 minutes prior to testing. Behaviors in *Mef2c*-Het mice were compared to control littermates tested on the same day. All behavioral tests were conducted using young adult mice (8-12 weeks), except juvenile communication (USV) recordings. Both male and female animals were included, except for adult ultrasonic vocalizations recordings. All behavior tests were conducted during the dark-phase (active-phase), and the experimenters were blind to genotypes.

#### Behavior Data Analysis

All data are presented as mean ± SEM. All comparisons were between littermates using appropriate two-sided statistical tests (specified in figure legends). Normal distribution of the data was assumed. Outliers were determined using GraphPad’s outlier calculator (Grubb’s) and excluded from analysis. *P*-values were calculated with unpaired *t*-test (two-tailed) or two-way ANOVA followed with Sidak’s multiple comparisons post-hoc test using GraphPad Prism, with specific tests described in figure legends.

#### Social Interaction

Mice were acclimated to the 3-arena sociability apparatus (Stoelting #60450) for 10 minutes. After acclimation, test mice were removed from the arena, and a novel, conspecific mouse and a novel object (medium black paper binder) were placed in a holding chamber within the side arenas. Test mice were returned to the arena and recorded for 10 minutes using ANY-maze behavior tracking software (Stoelting). Time is defined as time spent in each arena.

#### Ultrasonic vocalization recordings

Social ultrasonic vocalizations (USVs) were recorded from adult mice as previously described(Ey et al., 2013; Harrington et al., 2016). Briefly, ovariectomized female mice (C57BL/6; Jackson Labs) were injected with 15-μg estradiol 48 hours prior to testing and 1-mg progesterone 4 hours before testing to induce estrous. Test mice (8-12 week old male mice) were acclimated to a clean home-cage in a sound attenuated chamber for 5 minutes. After acclimation, an estrous female was introduced into the holding chamber with the male test mouse, and USVs were recorded for 5 minutes using Avisoft UltraSoundGate equipment (UltraSoundGate 116Hb with Condenser Microphone CM16; Avisoft Bioacoustics, Germany). USVs were analyzed using Avisoft SASLab Pro (Avisoft Bioacoustics) using a 20 kHz cutoff. *<u>Distress USVs</u>* were recorded from juvenile mice (pups) as previously described(Ey et al., 2013; Harrington et al., 2016; Scattoni et al., 2008). Briefly, individual pups of both sexes were identified with long-lasting subcutaneous tattoos (green tattoo paste; Ketchum) on the paws on post-natal day 4 (P4). Pups of both sexes were recorded in a random order in a small, sound-attenuated chamber following separation from dam and littermates. USVs were recorded for 3 minutes on post-natal days 7 and 10. USVs were quantified using Avisoft SASLab Pro (Avisoft Bioacoustics, German) with experiments blind to genotype.

#### Rotarod test

Mice were placed on the roller in a Rotarod apparatus (Ugo Basile Apparatus #47600) for a 2-minute training session with a rotation of 4 rpm or 8 rpm and replaced if mouse falls during training session. On the next day, test mice were returned to the roller, and the speed steadily increases from 4-40 rpms over 5 minutes. The latency and rpm for the mouse to fall off the roller were recorded. Each animal receives four testing sessions.

#### Locomotor Activity

Test mice were placed inside the Open Field Activity, Infrared Photobeam Activity Test Chamber (Med Associates), where an array of photobeams measures the mouse’s locomotor activity and jumping. Activity was monitored in the dark for one hour, and data are presented as the total activity during the hour.

#### Elevated Plus Maze (EPM)

Mice were introduced in the center of the elevated plus maze (Stoelting #60140) in white light (100 lux) and recorded for 5 minutes using ANY-maze behavior tracking software (Stoelting) with center-point detection. Data are reported as the percent of time spent in the open areas.

#### Fear Conditioning Test

Fear conditioning was performed as previously described(Wehner and Radcliffe, 2004). Briefly, test mice were placed in a fear conditioning chamber (Med Associates) and allowed to explore the arena for 2 minutes, after which a loud auditory stimulus (30 secs; 90 dB) that co-terminates with a 2-second mild foot-shock (0.5 mA) was presented to the animal. The mice were exposed to 3 tone/shock pairing with a 1-minute interval separating each tones/shock. The next day, animals were returned to the chamber and behavior (freezing) in the context, in a new context, and with the audible tone played in a new context is recorded with a video-tracking system (Video Freeze V2.7; Med Associates). Data are presented as percent of time the mouse is immobile.

#### Barnes Maze

Barnes maze was conducted as previously described(Rosenfeld and Ferguson, 2014) with modifications. Briefly, on the initial trial, mice were acclimated to the escape chamber for 2 minutes before testing. During testing, mice were introduced to the center of the Barnes Maze (Stoelting #60170) in bright white light (250 or 450 lux) with 4 distinct spatial cues evenly distributed around the arena. Mice were allowed to freely explore the arena for 5 minutes or until the mice found and entered the escape chamber. Latency to escape was recorded for each trial. Mice failing to find the escape chamber at the end of the trial were guided to the escape hole. All mice were left in the escape chamber for 2 minutes before being returned to their home cage. Each mouse was tested twice a day, and data reflect the average latency to enter the escape hole of the 2 trials per day.

#### Y-maze

Mice were introduced into one arm of a Y-maze (Stoelting #60180) with minimal white light (30 lux), and the mice were video tracked using ANY-maze behavior tracking software (Stoelting) for 5 minutes. Correct alternations were considered when the mouse entered a series of 3 different arms without re-entering a previously explored arm.

#### Novel Object Recognition (NOR)

Mice were acclimated for 10 minutes to an open field arena (OF; 44cm^2^) in dim light (30 lux) the day before testing. On test day, mice were acclimated to the OF for 5 minutes before objects were introduced. Mice were presented with 2 identical objects located on opposite sides of the OF arena and allowed to explore the objects for 10 minutes. One initial object was replaced with a novel object, and the mice were allowed to explore the objects for 10 minutes. Mice were recorded and analyzed using ANY-maze behavior tracking software (Stoelting). Interactions were considered when the center of the mouse was within 8 cms from the center of the object.

#### Sucrose preference

Test mice were singly housed and provided 2 identical ball-bearing sipper-style bottles to drink. Mice were acclimated to the 2 bottles for 4 days, where both bottles contained water on days 1 and 3 or sucrose solution on days 2 and 4. On days 5-8, mice were presented with 2 bottles, one with water and one with sucrose (1% (w/v)). Daily, the consumption of water and sucrose/quinine was measured, and the bottle position was altered to avoid potential side bias(Renthal et al., 2007). Data is presented as (solution consumption – water consumption) / total consumption = Preference Index.

#### Sucrose Self-Administration (SSA)

Sucrose self-administration (SA) was performed as previously described(Taniguchi et al., 2017). Briefly, mice were introduced to an operant conditioning chamber (Med Associates) at the same time each day during the dark cycle (active-phase). Both a light above the active nose poke hole and the house light indicated that sucrose was available. After an active hole nose poke, the availability lights went off and an internal light in the nose poke hole was activated. Active nose pokes immediately delivered a sucrose pellet (15 mg; TestDiet) and was followed by a 10 s time-out period. Inactive hole nose pokes did not have any consequences. After 12 days of acquiring, the mice entered a 7-day abstinence phase in their home cages and were not exposed to the operant conditioning chamber. Following abstinence, the mice were placed back in the operant chamber for 2 hours, where the sucrose pellets and cues were not present. Following 7 days of extinction, mice were re-introduced to the operant chamber and the availability cues and reward delivery cue (but no sucrose reward) were presented (cue-induced reinstatement). The number of active and inactive hole nose pokes were recorded during each session. Discrimination ratio is reported as number of active port entries / total number of port entries (active + inactive).

#### Shock and Acoustic Startle Response

Shock and acoustic startle response was performed as previously described(Harrington et al., 2016). Briefly, shock and startle responses were measured using Startle Reflex System and Advanced Startle software program (Med Associates). Mice were placed in Plexiglas and wire grid animal holders (ENV-264C) attached to a load cell platform (PHM-250) contained within a sound-attenuated chamber. For shock sensitivity, foot shocks (0.1 – 0.5 mA) were delivered by S/A Aversive Stimulators (ENV-414S) connected to the wire grid floors of the animal holder. For acoustic startle, mice were exposed to 5 white-noise pulses/amplitude (38ms) at 3 different amplitudes (70, 80, 90 dB) in a randomized order with variable inter-trial intervals (10-20s). Displacements of the load cell stabilimeter were converted into arbitrary units by an analog-to-digital converter interfaced to a personal computer.

#### RNA Isolation and Reverse Transcription PCR

Cortical tissue from p35-p40 mice (2 males and 2 females per genotype) were rapidly dissected and frozen at -80°C. Samples were thawed in TRIzol (Invitrogen), homogenized, and processed using the miRNeasy Mini Kit (Qiagen) according to manufacturer’s protocol. Total RNA was reverse-transcribed using Superscript III (Invitrogen) with random hexamers following manufacturer’s protocol. Quantitative real-time PCR was performed by the CFX96 qPCR instrument (Bio-Rad) using iTaq Universal SYBR Green Supermix (Bio-Rad) and primers specific to each target gene (Table S3). GAPDH was used to normalize gene expression in each sample.

#### RNA Sequencing

Total RNA was isolated from whole cortex of p35-p40 mice as described above. Sequencing was performed by BGI Americas Corporation (Cambridge, MA) using polyA mRNA isolation, directional RNA-seq library preparation, and the BGISEQ-500 platform with 150bp paired-end reads using DNA Nanoball (DNB) technology.

#### RNA-seq mapping, QC and expression quantification

Reads were aligned to the mouse mm10 reference genome using STAR 2.7.2a (Dobin et al., 2013) with the following parameters: “*-- outFilterMultimapNmax 10 --alignSJoverhangMin 10 --alignSJDBoverhangMin 1 -- outFilterMismatchNmax 3 --twopassMode Basic*”. For each sample, a BAM file including mapped and unmapped reads that spanned splice junctions was produced. Secondary alignment and multi-mapped reads were further removed using in-house scripts. Only uniquely mapped reads were retained for further analyses. Quality control metrics were performed using RseqQC using the m10 gene model provided. These steps include: number of reads after multiple-step filtering, ribosomal RNA reads depletion, and defining reads mapped to exons, UTRs, and intronic regions. Picard tool was implemented to refine the QC metrics (http://broadinstitute.github.io/picard/). Genecode annotation for mm10 (version M21) was used as reference alignment annotation and downstream quantification. Gene level expression was calculated using HTseq version 0.9.1 using intersection-strict mode by gene (Anders et al., 2015). Counts were calculated based on protein-coding genes from the annotation file.

#### Differential Expression

Counts were normalized using counts per million reads (CPM). Genes with no reads in either *Mef2c*-Het or WT samples were removed. Differential expression analysis was performed in R using linear modeling as following: *lm(gene expression ∼ Treatment)*

Fitting this model, we estimated log_2_ fold changes and P-values. P-values were adjusted for multiple comparisons using a Benjamini-Hochberg correction (FDR). Differentially expressed genes where consider for FDR < 0.05.

#### Functional Enrichment

The functional annotation of differentially expressed and co-expressed genes was performed using ToppGene (Chen et al., 2009). We used GO and KEGG databases. Pathways containing between 5 and 2000 genes were retained. A Benjamini-Hochberg FDR (*P* < 0.05) was applied as a multiple comparisons adjustment.

#### Gene set enrichment

Gene set enrichment was performed using a Fisher’s exact test in R with the following parameters: alternative = “greater”, conf.level = 0.95. We reported Odds Ratios (OR) and Benjamini-Hochberg adjusted P-value (FDR).

#### Availability of data and material

The NCBI Gene Expression Omnibus (GEO) accession number will be made available upon publication.

#### Code availability

Custom R codes and data to support the analysis, visualizations, functional and gene set enrichments will be made available upon publication.

#### Barrel Cortex Immunohistochemistry

Mice were transcardially perfused with 1X PBS followed by 4% paraformaldehyde (PFA) in PBS. Brains were hemisected and cortex was isolated. Cortices were flattened between 2 glass slides and post-fixed in 4% PFA for >24 hours. Cortices were cryoprotected in sucrose then sliced on a sliding stage microtome at 40 μM. Slices were blocked with 5% normal donkey serum and 1% albumin from bovine serum (0.3% TritonX-100, 1X PBS) for 2 hours, incubated with primary antibody anti-vGLUT2 (Abcam ab79157; 1:500) for overnight, incubated with Cy3 conjugated secondary antibody, and dehydrated. Cover slips were mounted using DPX mountant (Sigma).

#### Electrophysiology

All acute-slice electrophysiological experiments were performed in control and *Mef2c*-het mice at ages P30-P40. Acute coronal slices (300-µm thickness) containing barrel cortex were prepared in a semi-frozen 300 mOsM dissection solution containing (in mM): 100.0 choline chloride, 2.5 KCl, 1.25 Na2H2PO4, 25.0 NaHCO3, 25.0 D-glucose, 3.1 Na pyruvate, 9 Na ascorbate, 7.0 MgCl2, 0.5 CaCl2 and 5.0 kynurenic acid and was continually equilibrated with 95% O2 and 5% CO2 prior to and during the slicing procedure. Slices were transferred to a 315 mOsM normal artificial cerebrospinal fluid (ACSF) solution containing (in mM): 127 NaCl, 2.5 KCl, 1.20 Na2H2PO4, 24 NaHCO3, 11 D-glucose, 1.20 MgCl2, and 2 2.40 CaCl2, 0.4 Na Ascorbate to recover at 37°C for 30 minutes, and then transferred to room temperature ACSF for an additional 30 minutes prior to recording.

Layer 2/3 (L2/3) pyramidal neurons (depth 30-100 µm into the slice) of barrel cortex were visualized with infrared differential interference contrast optics (DIC/infrared optics) and identified by their location, apical dendrites, and burst spiking patterns in response to depolarizing current injection. Unless stated otherwise, all electrophysiological experiments were performed in whole cell voltage clamp mode at -70 mV using borosilicate pipettes (4-6 MΩ) made on NARISHIGE puller (NARISHIGE, PG10) from borosilicate tubing (Sutter Instruments) and filled by an internal solution containing (in mM): 120 K-Gluconate, 5 NaCl, 10 HEPES, 1.1 EGTA, 4 MgATP, 0.4 Na2GTP, 15 phosphocreatine, 2 MgCl2, and 0.1 CaCl2.

All data (Recordings) were acquired and analyzed by amplifier AXOPATCH 200B (Axon Instruments), digitizer BNC2090 (X National instruments) and software AxoGraph v.1.7.0, Clampfit v 8.0 (pClamp, Molecular devices) and Mini Analysis Program v.6.0.9 (Synaptosoft). Data were filtered at 2 kHz by AXOPATCH 200B amplifier (Axon Instruments) and digitized at 10-20 kHz via AxoGraph v.1.7.0.

#### Evoked postsynaptic currents

The evoked postsynaptic responses of BC pyramidal neurons in L2/3 were elicited by field stimulation of excitatory afferents in L4 directly underneath the recorded cells or in L2/3 of the adjustment barrel column at frequency of 0.05 Hz (0.05 c^-1^) - 3 stimulus in one second. The low-intensity pulses of stimulated current (25–100 mkA, 50–100 mks duration) were applied through a fine-tipped (∼2 mkm), bipolar stimulating electrode made from the borosilicate theta glass capillary tubing.

AMPA-receptor-mediated excitatory postsynaptic currents (EPSCs) were recorded in presence of picrotoxin (100 mkM, Sigma Aldrich) to block GABA_A_Rs. Inhibitory postsynaptic currents (IPSCs) mediated by GABA_A_ receptors were recorded in presence of DNQX (20 mkM, Tocris) to eliminate the current through AMPA-receptors.

For Paired-Pulse Ratio (PPR) measurements, two EPSC amplitudes were generated at -70mV with the inter-stimulus interval of 50 msec. The peak amplitude of the second EPSC (P2) was divided by the peak of the first amplitude (P1) to generate the PPR ratio (P2/P1).

#### Miniature postsynaptic current

Excitatory (E) and inhibitory (I) miniature postsynaptic currents (mPSCs) were recorded from L2/3 pyramidal neurons in voltage clamp mode at -70 mV. Pyramidal neurons were identified by their morphology parameters (pyramidal shape of soma, apical dendrite) and by bursting pattern of action potentials firing in response to depolarizing current injection. After identification, the normal ACSF was replaced by solution for mPSCs-records. For mEPSCs mediated by AMPARs, the extracellular bath solution (ACSF) contained 1 µM tetrodotoxin (TTX, Sigma Aldrich) and 100 µM pictrotoxin (Sigma Aldrich). mIPSCs were recorded using a high-chloride internal solution with a reversal potential at ECl- = -15 mV for chloride containing (in mM): 79 (70) K-gluconate, 44 (75) KCl, 6(2) NaCl, 10(11) HEPES, 0.2 EGTA, 4(2) MgATP, 0.4 (0.2) Na2GTP, 2 (1) MgCl2, and 0.1 CaCl2. For record GABA_A_Rs-mediated mIPSC the extracellular bath solution (ACSF) contained 1 µM TTX and 20 µM DNQX (AMPA-receptor antagonist, Sigma-Aldrich). For recording AMPARs-mediated mEPSC the extracellular bath solution (ACSF) contained 1 µM TTX and 100 µM picrotoxin (GABA_A_Rs antagonist, Sigma-Aldrich).

At the beginning of each sweep, a depolarizing step (4 mV for 100 ms) was generated to monitor series (10-40 MΩ) and input resistance (>400 MΩ). Data were collected in a series of traces until >300 events were recorded. Synaptic events were detected via custom parameters in MiniAnalysis software (Synaptosoft, Decatur, GA) and subsequently confirmed by observer. For each event, amplitude and frequency was measured and used to determine average mean.

#### Dendritic Spine Analysis

Dendritic spine labeling was done as previously described(Spencer et al., 2018) with some modifications. Briefly, mice were lightly transcardially perfused with 1.5% PFA in PBS and brains were post-fixed for 1 hour at 4°C in 1.5% PFA. Brains were sectioned at 200 μm using a vibrating microtome (Leica VT100P). Tungsten particles (1.3 μm diameter; Bio-Rad) were coated with lipophilic carbocyanine dye DiI (Life Technologies) and diolistically delivered into cortical regions using a Helios Gene Gun system (Bio-Rad) fitted with a polycarbonate filter (3.0 μm pore size; BD Biosciences). After delivering the DiI-coated particles, slices were incubated in PBS at 4°C overnight to allow the dye to diffuse along the neuronal dendrites and axons, then post-fixed for 1 hour in 4% PFA before mounting with Prolong^®^ Gold Antifade (Invitrogen) and imaging. Images of secondary and tertiary L2/3 apical (4-7 segments per animal, 31 segments per group) and tertiary basal dendrites (3-7 segments per animal, WT=31 segments, het=26 segments) and L5 tertiary basal dendrites (3-5 segments per animal, WT=23 segments, het=21 segments) in the barrel field were imaged with a Leica SP8 laser scanning confocal microscope equipped with HyD detectors for improved sensitivity. 50 µm dendritic segments were imaged with a 63X oil immersion objective (1.4 NA) at 1024x512 frame size, 4.1x digital zoom, and 0.1 µm Z-step size, generating a voxel size of 44 x 44 x 100 nm. DiI was excited using an OPSL 552 laser line with a pinhole of 0.8 Airy Units (AU). Laser power and gain were initially optimized, and then held relatively constant for the remainder of the experiment (i.e. laser power and gain were dynamically adjusted to avoid saturated voxels). Imaging parameters were chosen based off the Nyquist theorem as recommended by Huygens software (Scientific Volume Imaging, Hilversum, NL), which was used for deconvolution. Deconvolved images were imported into BitPlane Imaris (Version 9.1, Zurich, CH) for 3D reconstruction. The filament module was used to trace each dendrite, and the autopath tool was used to assign spines across each dendrite. Spine density per µm of dendrite and average spine head diameter of each dendritic segment were exported as variables.

#### Tamoxifen Treatment of *Mef2c cHet^Cx3Cr1^* and *Cx3Cr1^creER/+^*; Ai14 Mice

To induce recombination of the *Mef2c* floxed allele or Gt(ROSA)26Sor locus to express tdTomato in microglia, *Cx3Cr1^creER/+^*, *Mef2c^fl/+^* (experimental) and *Cx3Cr1^creER/+^*, *Mef2c^+/+^* (control) or *Cx3Cr1^creER/+^*; Ai14 pups were treated with tamoxifen via mother’s milk (100 mg/kg of tamoxifen in 10% ethanol/90% sesame oil i.p. injection of dam) from post-natal day 1 through post-natal day 3. Starting at post-natal day 4, treated pups were then fostered by a lactating CD-1 female mouse until weaning.

#### Immunohistochemistry

Mice were terminally anesthetized with ketamine/xylazine and transcardially perfused with PBS followed by 4% (w/v) paraformaldehyde (PFA). Brains were post-fixed for >24 hours at 4°C in 4% PFA then cryoprotected in 30% sucrose. Brains were coronally sectioned at 40-50 μm using a sliding microtome and stored in 1X PBS with 0.02% sodium azide. Anti-IBA1 IHC was performed based on manufacturer’s protocol with modifications (Wako). Sections were washed 3 times for 10 minutes with 1 X PBS with TritonX-100 (0.3%). Sections were then blocked with 1X PBS with 1% BSA and 0.3% TritonX-100 for 2 hours. Sections were immunostained with primary rabbit anti-IBA1 antibody (1:1000; Wako 019-19741) overnight at 4°C followed by Cy3 conjugated goat anti-rabbit secondary antibody (1:400 for 2 hours) or Alexa-Fluor-647 donkey anti-rabbit secondary antibody (1:400 for 2 hours). Sections were mounted with Prolong Gold with DAPI mountant (ThermoFisher).

Anti-GFP and anti-MEF2C IHC was performed as described in the following. Sections were washed 3 times for 10 minutes with 1 X PBS with TritonX-100 (0.3%) and then blocked in 3% bovine serum albumin, 3% normal donkey serum, 0.3% triton X-100, 0.2% tween-20 in 1X PBS. Sections were immunostained with primary chicken anti-GFP (1:1000; Aves GFP-1020) antibody and rabbit anti-MEF2A/MEF2C (1:300; abcam ab197070) antibody overnight at 4°C followed by Alexa Fluor-488 donkey anti-chicken (1:400) and Cy3 donkey anti-rabbit secondary antibody (1:200) for 1.5 hours. Sections were mounted with Prolong Gold with DAPI mountant (ThermoFisher).

#### Confocal Imaging and Image Analysis

Iba1 stained sections were imaged on a Zeiss 880 confocal microscope at 20x objective. On average, 3 images were imaged per mouse/brain region. Images were deconvolved in AutoQuant (Bitplane) and analyzed in Imaris software (Bitplane). Imaris analysis began with surface rendering of Iba1+ cells. The mean intensity of each cell, cell number, and cell soma volume were calculated in the Imaris software. The mean intensity of each cell by animal was entered into GraphPad Prism software to generate cumulative frequency distributions. Statistical testing was performed on this data via Kolmogorov–Smirnov test. Iba1+ cell numbers per field and cell soma volumes were entered into GraphPad Prism by genotype and brain region. Genotype-based differences in Iba1+ cell numbers were assessed via unpaired two-tailed or nested t-tests.

Imaging of MEF2C expression in microglia was performed on a Zeiss 880 confocal microscope at 40x objective.

Imaging to confirm microglia-specific recombination was performed in *Cx3Cr1^creER/+^*; Ai14 mice that express tdTomato under the Gt(ROSA)26Sor locus. Stained sections were imaged on a Zeiss 880 confocal microscope at 20x objective. Quantification was performed in Fiji by counting.

#### Cytokine Antibody Array

The cytokine antibody array was performed according to manufacturer’s instructions except the membranes were blocked for 2 hours. A starting concentration of 450 μg of cortical brain tissue from Mef2c-Het and controls were used for each membrane.

## References

Adachi, M., Lin, P.Y., Pranav, H., and Monteggia, L.M. (2015). Postnatal Loss of Mef2c Results in Dissociation of Effects on Synapse Number and Learning and Memory. Biol Psychiatry.

Anders, S., Pyl, P.T., and Huber, W. (2015). HTSeq--a Python framework to work with high-throughput sequencing data. Bioinformatics 31, 166–169.

Antoine, M.W., Langberg, T., Schnepel, P., and Feldman, D.E. (2019). Increased Excitation-Inhibition Ratio Stabilizes Synapse and Circuit Excitability in Four Autism Mouse Models. Neuron 101, 648–661 e644.

Arnold, M.A., Kim, Y., Czubryt, M.P., Phan, D., McAnally, J., Qi, X., Shelton, J.M., Richardson, J.A., Bassel-Duby, R., and Olson, E.N. (2007). MEF2C transcription factor controls chondrocyte hypertrophy and bone development. Developmental cell 12, 377–389.

Assali, A., Harrington, A.J., and Cowan, C.W. (2019). Emerging roles for MEF2 in brain development and mental disorders. Current opinion in neurobiology 59, 49–58.

Barbosa, A.C., Kim, M.S., Ertunc, M., Adachi, M., Nelson, E.D., McAnally, J., Richardson, J.A., Kavalali, E.T., Monteggia, L.M., Bassel-Duby, R., and Olson, E.N. (2008). MEF2C, a transcription factor that facilitates learning and memory by negative regulation of synapse numbers and function. Proceedings of the National Academy of Sciences of the United States of America 105, 9391–9396.

Barski, J.J., Dethleffsen, K., and Meyer, M. (2000). Cre recombinase expression in cerebellar Purkinje cells. Genesis 28, 93–98.

Berland, S., and Houge, G. (2010). Late-onset gain of skills and peculiar jugular pit in an 11-year-old girl with 5q14.3 microdeletion including MEF2C. Clin Dysmorphol 19, 222–224.

Bialas, A.R., and Stevens, B. (2013). TGF-β signaling regulates neuronal C1q expression and developmental synaptic refinement. Nature Neuroscience 16, 1773.

Bienvenu, T., Diebold, B., Chelly, J., and Isidor, B. (2013). Refining the phenotype associated with MEF2C point mutations. Neurogenetics 14, 71–75.

Chen, J., Bardes, E.E., Aronow, B.J., and Jegga, A.G. (2009). ToppGene Suite for gene list enrichment analysis and candidate gene prioritization. Nucleic acids research 37, W305–311.

Deczkowska, A., Matcovitch-Natan, O., Tsitsou-Kampeli, A., Ben-Hamo, S., Dvir-Szternfeld, R., Spinrad, A., Singer, O., David, E., Winter, D.R., Smith, L.K., et al. (2017). Mef2C restrains microglial inflammatory response and is lost in brain ageing in an IFN-I-dependent manner. Nature communications 8, 717.

Derecki, N.C., Cronk, J.C., Lu, Z., Xu, E., Abbott, S.B., Guyenet, P.G., and Kipnis, J. (2012). Wild-type microglia arrest pathology in a mouse model of Rett syndrome. Nature 484, 105–109.

Dobin, A., Davis, C.A., Schlesinger, F., Drenkow, J., Zaleski, C., Jha, S., Batut, P., Chaisson, M., and Gingeras, T.R. (2013). STAR: ultrafast universal RNA-seq aligner. Bioinformatics 29, 15–21.

Engels, H., Wohlleber, E., Zink, A., Hoyer, J., Ludwig, K.U., Brockschmidt, F.F., Wieczorek, D., Moog, U., Hellmann-Mersch, B., Weber, R.G., et al. (2009). A novel microdeletion syndrome involving 5q14.3-q15: clinical and molecular cytogenetic characterization of three patients. European journal of human genetics: EJHG 17, 1592–1599.

Ey, E., Torquet, N., Le Sourd, A.M., Leblond, C.S., Boeckers, T.M., Faure, P., and Bourgeron, T. (2013). The Autism ProSAP1/Shank2 mouse model displays quantitative and structural abnormalities in ultrasonic vocalisations. Behavioural brain research 256, 677–689.

Fioravante, D., and Regehr, W.G. (2011). Short-term forms of presynaptic plasticity. Current opinion in neurobiology 21, 269–274.

Flavell, S.W., Cowan, C.W., Kim, T.K., Greer, P.L., Lin, Y., Paradis, S., Griffith, E.C., Hu, L.S., Chen, C., and Greenberg, M.E. (2006). Activity-dependent regulation of MEF2 transcription factors suppresses excitatory synapse number. Science 311, 1008–1012.

Flavell, S.W., Kim, T.K., Gray, J.M., Harmin, D.A., Hemberg, M., Hong, E.J., Markenscoff-Papadimitriou, E., Bear, D.M., and Greenberg, M.E. (2008). Genome-wide analysis of MEF2 transcriptional program reveals synaptic target genes and neuronal activity-dependent polyadenylation site selection. Neuron 60, 1022–1038.

Gandal, M.J., Zhang, P., Hadjimichael, E., Walker, R.L., Chen, C., Liu, S., Won, H., van Bakel, H., Varghese, M., Wang, Y., et al. (2018). Transcriptome-wide isoform-level dysregulation in ASD, schizophrenia, and bipolar disorder. Science 362.

Garber, K. (2007). Neuroscience. Autism’s cause may reside in abnormalities at the synapse. Science 317, 190–191.

Gosselin, D., Skola, D., Coufal, N.G., Holtman, I.R., Schlachetzki, J.C.M., Sajti, E., Jaeger, B.N., O’Connor, C., Fitzpatrick, C., Pasillas, M.P., et al. (2017). An environment-dependent transcriptional network specifies human microglia identity. Science 356.

Hagemeyer, N., Hanft, K.-M., Akriditou, M.-A., Unger, N., Park, E.S., Stanley, E.R., Staszewski, O., Dimou, L., and Prinz, M. (2017). Microglia contribute to normal myelinogenesis and to oligodendrocyte progenitor maintenance during adulthood. Acta Neuropathol 134, 441–458.

Harrington, A.J., Raissi, A., Rajkovich, K., Berto, S., Kumar, J., Molinaro, G., Raduazzo, J., Guo, Y., Loerwald, K., Konopka, G., et al. (2016). MEF2C regulates cortical inhibitory and excitatory synapses and behaviors relevant to neurodevelopmental disorders. Elife 5.

Hoogland, I.C., Houbolt, C., van Westerloo, D.J., van Gool, W.A., and van de Beek, D. (2015). Systemic inflammation and microglial activation: systematic review of animal experiments. J Neuroinflammation 12, 114.

Horiuchi, M., Smith, L., Maezawa, I., and Jin, L.W. (2017). CX3CR1 ablation ameliorates motor and respiratory dysfunctions and improves survival of a Rett syndrome mouse model. Brain Behav Immun 60, 106–116.

Ito, D., Imai, Y., Ohsawa, K., Nakajima, K., Fukuuchi, Y., and Kohsaka, S. (1998). Microglia-specific localisation of a novel calcium binding protein, Iba1. Molecular Brain Research 57, 1-9.

Ito, D., Tanaka, K., Suzuki, S., Dembo, T., and Fukuuchi, Y. (2001). Enhanced Expression of Iba1, Ionized Calcium-Binding Adapter Molecule 1, After Transient Focal Cerebral Ischemia In Rat Brain. Stroke 32, 1208–1215.

Iwasato, T., Inan, M., Kanki, H., Erzurumlu, R.S., Itohara, S., and Crair, M.C. (2008). Cortical adenylyl cyclase 1 is required for thalamocortical synapse maturation and aspects of layer IV barrel development. The Journal of neuroscience: the official journal of the Society for Neuroscience 28, 5931–5943.

Kamath, S.P., and Chen, A.I. (2018). Myocyte Enhancer Factor 2c Regulates Dendritic Complexity and Connectivity of Cerebellar Purkinje Cells. Mol Neurobiol.

Kang, J., Gocke, C.B., and Yu, H. (2006). Phosphorylation-facilitated sumoylation of MEF2C negatively regulates its transcriptional activity. BMC Biochem 7, 5.

Le Meur, N., Holder-Espinasse, M., Jaillard, S., Goldenberg, A., Joriot, S., Amati-Bonneau, P., Guichet, A., Barth, M., Charollais, A., Journel, H., et al. (2010). MEF2C haploinsufficiency caused by either microdeletion of the 5q14.3 region or mutation is responsible for severe mental retardation with stereotypic movements, epilepsy and/or cerebral malformations. Journal of medical genetics 47, 22–29.

Li, H., Radford, J.C., Ragusa, M.J., Shea, K.L., McKercher, S.R., Zaremba, J.D., Soussou, W., Nie, Z., Kang, Y.J., Nakanishi, N., et al. (2008). Transcription factor MEF2C influences neural stem/progenitor cell differentiation and maturation in vivo. Proceedings of the National Academy of Sciences of the United States of America 105, 9397–9402.

Li, Q., and Barres, B.A. (2018). Microglia and macrophages in brain homeostasis and disease. Nat Rev Immunol 18, 225–242.

Lyons, M.R., Schwarz, C.M., and West, A.E. (2012). Members of the myocyte enhancer factor 2 transcription factor family differentially regulate Bdnf transcription in response to neuronal depolarization. The Journal of neuroscience: the official journal of the Society for Neuroscience 32, 12780–12785.

Mayer, C., Hafemeister, C., Bandler, R.C., Machold, R., Batista Brito, R., Jaglin, X., Allaway, K., Butler, A., Fishell, G., and Satija, R. (2018). Developmental diversification of cortical inhibitory interneurons. Nature 555, 457–462.

McKinsey, T.A., Zhang, C.L., and Olson, E.N. (2002). MEF2: a calcium-dependent regulator of cell division, differentiation and death. Trends in biochemical sciences 27, 40–47.

Mikhail, F.M., Lose, E.J., Robin, N.H., Descartes, M.D., Rutledge, K.D., Rutledge, S.L., Korf, B.R., and Carroll, A.J. (2011). Clinically relevant single gene or intragenic deletions encompassing critical neurodevelopmental genes in patients with developmental delay, mental retardation, and/or autism spectrum disorders. American journal of medical genetics Part A 155A, 2386-2396.

Morrow, E.M., Yoo, S.Y., Flavell, S.W., Kim, T.K., Lin, Y., Hill, R.S., Mukaddes, N.M., Balkhy, S., Gascon, G., Hashmi, A., et al. (2008). Identifying autism loci and genes by tracing recent shared ancestry. Science 321, 218–223.

Novara, F., Beri, S., Giorda, R., Ortibus, E., Nageshappa, S., Darra, F., Dalla Bernardina, B., Zuffardi, O., and Van Esch, H. (2010). Refining the phenotype associated with MEF2C haploinsufficiency. Clinical genetics 78, 471–477.

Odell, D., Maciulis, A., Cutler, A., Warren, L., McMahon, W.M., Coon, H., Stubbs, G., Henley, K., and Torres, A. (2005). Confirmation of the association of the C4B null allelle in autism. Human Immunology 66, 140–145.

Paciorkowski, A.R., Traylor, R.N., Rosenfeld, J.A., Hoover, J.M., Harris, C.J., Winter, S., Lacassie, Y., Bialer, M., Lamb, A.N., Schultz, R.A., et al. (2013). MEF2C Haploinsufficiency features consistent hyperkinesis, variable epilepsy, and has a role in dorsal and ventral neuronal developmental pathways. Neurogenetics 14, 99–111.

Paolicelli, R.C., Bolasco, G., Pagani, F., Maggi, L., Scianni, M., Panzanelli, P., Giustetto, M., Ferreira, T.A., Guiducci, E., Dumas, L., et al. (2011). Synaptic Pruning by Microglia Is Necessary for Normal Brain Development. Science 333, 1456.

Parkhurst, C.N., Yang, G., Ninan, I., Savas, J.N., Yates, J.R., 3rd, Lafaille, J.J., Hempstead, B.L., Littman, D.R., and Gan, W.B. (2013). Microglia promote learning-dependent synapse formation through brain-derived neurotrophic factor. Cell 155, 1596-1609.

Pfeiffer, B.E., Zang, T., Wilkerson, J.R., Taniguchi, M., Maksimova, M.A., Smith, L.N., Cowan, C.W., and Huber, K.M. (2010). Fragile X mental retardation protein is required for synapse elimination by the activity-dependent transcription factor MEF2. Neuron 66, 191–197.

Pont-Lezica, L., Beumer, W., Colasse, S., Drexhage, H., Versnel, M., and Bessis, A. (2014). Microglia shape corpus callosum axon tract fasciculation: functional impact of prenatal inflammation. Eur J Neurosci 39, 1551–1557.

Pulipparacharuvil, S., Renthal, W., Hale, C.F., Taniguchi, M., Xiao, G., Kumar, A., Russo, S.J., Sikder, D., Dewey, C.M., Davis, M.M., et al. (2008). Cocaine regulates MEF2 to control synaptic and behavioral plasticity. Neuron 59, 621–633.

Rajkovich, K.E., Loerwald, K.W., Hale, C.F., Hess, C.T., Gibson, J.R., and Huber, K.M. (2017). Experience-Dependent and Differential Regulation of Local and Long-Range Excitatory Neocortical Circuits by Postsynaptic Mef2c. Neuron 93, 48–56.

Renthal, W., Maze, I., Krishnan, V., Covington, H.E., 3rd, Xiao, G., Kumar, A., Russo, S.J., Graham, A., Tsankova, N., Kippin, T.E., et al. (2007). Histone deacetylase 5 epigenetically controls behavioral adaptations to chronic emotional stimuli. Neuron 56, 517-529.

Rosenfeld, C.S., and Ferguson, S.A. (2014). Barnes maze testing strategies with small and large rodent models. J Vis Exp, e51194.

Saunders, A., Macosko, E.Z., Wysoker, A., Goldman, M., Krienen, F.M., de Rivera, H., Bien, E., Baum, M., Bortolin, L., Wang, S., et al. (2018). Molecular Diversity and Specializations among the Cells of the Adult Mouse Brain. Cell 174, 1015–1030 e1016.

Scattoni, M.L., Gandhy, S.U., Ricceri, L., and Crawley, J.N. (2008). Unusual repertoire of vocalizations in the BTBR T+tf/J mouse model of autism. PloS one 3, e3067.

Schafer, D.P., Heller, C.T., Gunner, G., Heller, M., Gordon, C., Hammond, T., Wolf, Y., Jung, S., and Stevens, B. (2016). Microglia contribute to circuit defects in Mecp2 null mice independent of microglia-specific loss of Mecp2 expression. Elife 5.

Schafer, D.P., Lehrman, E.K., Kautzman, A.G., Koyama, R., Mardinly, A.R., Yamasaki, R., Ransohoff, R.M., Greenberg, M.E., Barres, B.A., and Stevens, B. (2012). Microglia sculpt postnatal neural circuits in an activity and complement-dependent manner. Neuron 74, 691–705.

Sekar, A., Bialas, A.R., de Rivera, H., Davis, A., Hammond, T.R., Kamitaki, N., Tooley, K., Presumey, J., Baum, M., Van Doren, V., et al. (2016). Schizophrenia risk from complex variation of complement component 4. Nature 530, 177.

Shigemoto-Mogami, Y., Hoshikawa, K., Goldman, J.E., Sekino, Y., and Sato, K. (2014). Microglia Enhance Neurogenesis and Oligodendrogenesis in the Early Postnatal Subventricular Zone. The Journal of Neuroscience 34, 2231.

Sierra, A., Encinas, J.M., Deudero, J.J.P., Chancey, J.H., Enikolopov, G., Overstreet-Wadiche, L.S., Tsirka, S.E., and Maletic-Savatic, M. (2010). Microglia Shape Adult Hippocampal Neurogenesis through Apoptosis-Coupled Phagocytosis. Cell Stem Cell 7, 483–495.

Spencer, S., Neuhofer, D., Chioma, V.C., Garcia-Keller, C., Schwartz, D.J., Allen, N., Scofield, M.D., Ortiz-Ithier, T., and Kalivas, P.W. (2018). A Model of Delta(9)-Tetrahydrocannabinol Self-administration and Reinstatement That Alters Synaptic Plasticity in Nucleus Accumbens. Biol Psychiatry 84, 601–610.

Stevens, B., Allen, N.J., Vazquez, L.E., Howell, G.R., Christopherson, K.S., Nouri, N., Micheva, K.D., Mehalow, A.K., Huberman, A.D., Stafford, B., et al. (2007). The classical complement cascade mediates CNS synapse elimination. Cell 131, 1164–1178.

Taniguchi, M., Carreira, M.B., Cooper, Y.A., Bobadilla, A.C., Heinsbroek, J.A., Koike, N., Larson, E.B., Balmuth, E.A., Hughes, B.W., Penrod, R.D., et al. (2017). HDAC5 and Its Target Gene, Npas4, Function in the Nucleus Accumbens to Regulate Cocaine-Conditioned Behaviors. Neuron 96, 130-144 e136.

Tonk, V., Kyhm, J.H., Gibson, C.E., and Wilson, G.N. (2011). Interstitial deletion 5q14.3q21.3 with MEF2C haploinsufficiency and mild phenotype: when more is less. American journal of medical genetics Part A 155A, 1437-1441.

Tsai, N.P., Wilkerson, J.R., Guo, W., Maksimova, M.A., DeMartino, G.N., Cowan, C.W., and Huber, K.M. (2012). Multiple autism-linked genes mediate synapse elimination via proteasomal degradation of a synaptic scaffold PSD-95. Cell 151, 1581–1594.

Tu, S., Akhtar, M.W., Escorihuela, R.M., Amador-Arjona, A., Swarup, V., Parker, J., Zaremba, J.D., Holland, T., Bansal, N., Holohan, D.R., et al. (2017). NitroSynapsin therapy for a mouse MEF2C haploinsufficiency model of human autism. Nature communications 8, 1488.

Velmeshev, D., Schirmer, L., Jung, D., Haeussler, M., Perez, Y., Mayer, S., Bhaduri, A., Goyal, N., Rowitch, D.H., and Kriegstein, A.R. (2019). Single-cell genomics identifies cell type-specific molecular changes in autism. Science 364, 685–689.

Vrecar, I., Innes, J., Jones, E.A., Kingston, H., Reardon, W., Kerr, B., Clayton-Smith, J., and Douzgou, S. (2017). Further Clinical Delineation of the MEF2C Haploinsufficiency Syndrome: Report on New Cases and Literature Review of Severe Neurodevelopmental Disorders Presenting with Seizures, Absent Speech, and Involuntary Movements. J Pediatr Genet 6, 129–141.

Wang DZ, Valdez MR, McAnally J, Richardson J, and EN., O. (2001). The Mef2c gene is a direct transcriptional target of myogenic bHLH and MEF2 proteins during skeletal muscle development. Development 128, 4623–4633.

Wang, J., Wegener, J.E., Huang, T.-W., Sripathy, S., De Jesus-Cortes, H., Xu, P., Tran, S., Knobbe, W., Leko, V., Britt, J., et al. (2015). Wild-type microglia do not reverse pathology in mouse models of Rett syndrome. Nature 521, E1.

Wehner, J.M., and Radcliffe, R.A. (2004). Cued and contextual fear conditioning in mice. Curr Protoc Neurosci Chapter 8, Unit 8 5C.

Wright-Jin, E.C., and Gutmann, D.H. (2019). Microglia as Dynamic Cellular Mediators of Brain Function. Trends Mol Med.

Zang, T., Maksimova, M.A., Cowan, C.W., Bassel-Duby, R., Olson, E.N., and Huber, K.M. (2013). Postsynaptic FMRP bidirectionally regulates excitatory synapses as a function of developmental age and MEF2 activity. Molecular and cellular neurosciences 56, 39–49.

Zhan, Y., Paolicelli, R.C., Sforazzini, F., Weinhard, L., Bolasco, G., Pagani, F., Vyssotski, A.L., Bifone, A., Gozzi, A., Ragozzino, D., and Gross, C.T. (2014). Deficient neuron-microglia signaling results in impaired functional brain connectivity and social behavior. Nat Neurosci 17, 400–406.

Zhang, Y., Chen, K., Sloan, S.A., Bennett, M.L., Scholze, A.R., Keeffe, S., Phatnani, H.P., Guarnieri, P., Caneda, C., Ruderisch, N., et al. (2014a). An RNA-Sequencing Transcriptome and Splicing Database of Glia, Neurons, and Vascular Cells of the Cerebral Cortex. The Journal of Neuroscience 34, 11929.

Zhang, Y., Chen, K., Sloan, S.A., Bennett, M.L., Scholze, A.R., O’Keeffe, S., Phatnani, H.P., Guarnieri, P., Caneda, C., Ruderisch, N., et al. (2014b). An RNA-sequencing transcriptome and splicing database of glia, neurons, and vascular cells of the cerebral cortex. The Journal of neuroscience: the official journal of the Society for Neuroscience 34, 11929–11947.

Zoghbi, H.Y., and Bear, M.F. (2012). Synaptic dysfunction in neurodevelopmental disorders associated with autism and intellectual disabilities. Cold Spring Harbor perspectives in biology 4.

Zweier, M., Gregor, A., Zweier, C., Engels, H., Sticht, H., Wohlleber, E., Bijlsma, E.K., Holder, S.E., Zenker, M., Rossier, E., et al. (2010). Mutations in MEF2C from the 5q14.3q15 microdeletion syndrome region are a frequent cause of severe mental retardation and diminish MECP2 and CDKL5 expression. Human mutation 31, 722–733.

Zweier, M., and Rauch, A. (2012). The MEF2C-Related and 5q14.3q15 Microdeletion Syndrome. Molecular syndromology 2, 164–170.

